# A multidimensional framework for measuring biotic novelty: How novel is a community?

**DOI:** 10.1101/824045

**Authors:** Conrad Schittko, Maud Bernard-Verdier, Tina Heger, Sascha Buchholz, Ingo Kowarik, Moritz von der Lippe, Birgit Seitz, Jasmin Joshi, Jonathan M. Jeschke

## Abstract

Anthropogenic changes in climate, land use and disturbance regimes, as well as introductions of non-native species can lead to the transformation of many ecosystems. The resulting novel ecosystems are usually characterized by species assemblages that have not occurred previously in a given area. Quantifying the ecological novelty of communities (i.e. biotic novelty) would enhance the understanding of environmental change. However, quantification remains challenging since current novelty metrics, such as the number and/or proportion of non-native species in a community, fall short of considering both functional and evolutionary aspects of biotic novelty. Here, we propose the Biotic Novelty Index (BNI), an intuitive and flexible multidimensional measure that combines (1) functional differences between native and non-native introduced species with (2) temporal dynamics of species introductions. We show that the BNI is an additive partition of Rao’s quadratic entropy, capturing the novel interaction component of the community’s functional diversity. Simulations show that the index varies predictably with the relative amount of functional novelty added by recently arrived species, and they illustrate the need to provide an additional standardized version of the index. We present a detailed R-code and two applications of the BNI by (1) measuring changes of biotic novelty of dry grassland plant communities along an urbanization gradient in a metropolitan region and (2) determining the biotic novelty of plant species assemblages at a national scale. Results illustrate the applicability of the index across scales and its flexibility in the use of data of different quality. Both case studies revealed strong connections between biotic novelty and increasing urbanization, a measure of abiotic novelty. We conclude that the BNI framework may help in building a basis for a better understanding of the ecological and evolutionary consequences of global change.

## Introduction

Ecological novelty has received growing attention in the recent literature (e.g. Hobbs *et al*. 2006; Heger *et al*. 2019) focusing on novel organisms (Jeschke *et al*. 2013), novel species interactions (Pearse & Altermatt 2013; Bezemer *et al*. 2014; Carthey & Banks 2014), novel communities (Lurgi *et al*. 2012) or novel ecosystems (Hobbs *et al*. 2009, 2013; Higgs 2017). One major aspect of ecological novelty is the emergence of abiotic and biotic conditions that are beyond the historical range of conditions at a given site or area (Mora *et al*. 2013), sometimes without present or past analog conditions anywhere else (Williams & Jackson 2007). A site can be novel in terms of abiotic conditions, resulting for example from changes in climate, nitrogen deposition or pollution by microplastics. Novelty can also result from changes in species composition, structure or ecological processes, generating biotic novelty (Heger *et al*. 2019). Furthermore, abiotic novelty can cause biotic novelty (Chapin & Starfield 1997; Williams & Jackson 2007; Bogan & Lytle 2011; Correa-Metrio *et al*. 2012), such as a when a reshuffling of species is induced by climate change (Williams & Jackson 2007) – and vice versa when introduced species, for example, strongly affect the nutrient cycling (Vilà *et al*. 2011; Jäger *et al*. 2013). At the same time, biotic novelty can occur without abiotic novelty: a non-native species introduction may create novelty in species composition, whereas abiotic conditions remain essentially unchanged. Hence, rigorously measuring novelty requires explicit definition of the relevant variables (Radeloff *et al*. 2015).

However, the question of how to quantify ecological novelty in a standardized and comparable manner has rarely been considered. A straightforward approach to measuring abiotic novelty is to compare current abiotic variables, for instance climatic variables, in an area with their historic values by applying dissimilarity metrics (Williams *et al*. 2007; Garcia *et al*. 2014; Radeloff *et al*. 2015). This approach has become increasingly common in climate change science, and may be applied to any abiotic factor for which reference data are available.

A common measure of biotic novelty is simply the number and/or proportion of novel species (e.g. non-native species) in a community (Parker *et al*. 2006; Qian & Ricklefs 2006; Wilsey *et al*. 2009; Catford *et al*. 2012; Korell *et al*. 2016). However, assigning species to one of these two categories is a broad generalization and temporal dynamics of novel species introductions and their interactions with native species are reduced to a binary view. In a given community, species usually differ in their residence time in the focal region, depending on the time of arrival mediated by natural or anthropogenic pathways (Fig. 1). This has evolutionary consequences since both native and non-native species may gradually adapt to their new interaction partner(s) over time (Strauss *et al*. 2006; Verhoeven *et al*. 2009; Carthey & Banks 2012; Hulme & Bernard-Verdier 2018), which may lead to a decrease of novelty in the community (Saul & Jeschke 2015). Consequently, we argue that a quantification method of biotic novelty should include a component that captures the different time spans of coexistence of the species in a given community.

**Figure 1:**
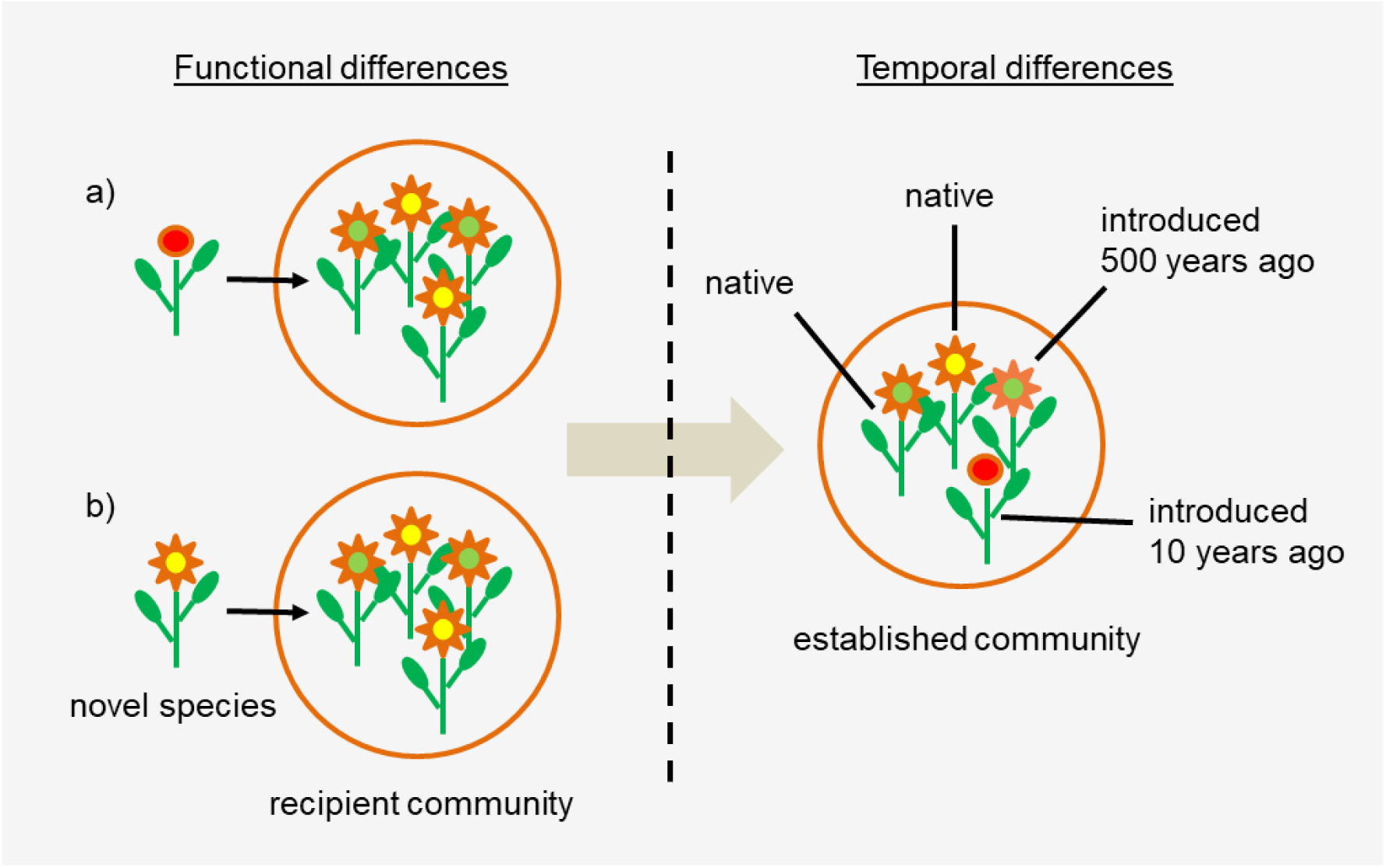
Scheme of two aspects of biotic novelty in a hypothetical plant community that are both captured by the BNI. Left side: A novel species that enters a community of resident species may be functionally different (scenario a) or similar (scenario b) compared to the resident species. Right side: In a given community, there is typically not only one nonresident species, but multiple species that may have arrived at different points in time in the focal region.

Another limitation of assessing biotic novelty only by quantifying native vs. non-native species is the omission of functional differences between species. A novel species that enters a community may be functionally similar or different compared with the resident species (Fig. 1; Hulme & Bernard-Verdier 2018). We argue that a species that is functionally dissimilar from the resident species represents greater biotic novelty than one that is similar to the pre-existing community.

Several recent studies proposed new approaches to capture the biotic novelty of ecological communities (Baselga 2010; Saul *et al*. 2013; Helm *et al*. 2015; Shimadzu *et al*. 2015). These approaches mainly focus on community dynamics and species turnover over time. For example, Shimadzu *et al*. (2015) converted commonly used measures of β-diversity, such as Jaccard’s index of dissimilarity, to a measure of temporal β-diversity that compares the species composition of one community at two points in time (i.e. at an initial state and the current state). This provides a powerful way to quantify novelty compared to past “reference states” (Heger *et al*. 2019), but it is not easily applicable to compare two existing communities for which local temporal dynamics data are missing.

We propose a new multidimensional measure of biotic novelty called Biotic Novelty Index (BNI), which serves to capture the two components of novelty as described by Heger *et al*. (2019): (1) a change-dependent (“different”) component and (2) a time-dependent (“before”) component. In this sense, a situation is ecologically novel if the new situation is “different”, e.g. in terms of species composition, from the situation that was present “before”, e.g. compared to historic baseline conditions. Accordingly, our index relies on: (1) pairwise dissimilarities between species (e.g. functional or phylogenetic distances), and (2) the residence time of each species in the area considered. The index was designed to make comparisons of novelty between several communities (e.g. along gradients) at the present point in time, without prior knowledge of the local communities assembly history. The BNI is based on the formula for Rao’s quadratic entropy (hereafter Rao’s Q; Rao 1982; Botta-Dukát 2005), which is one of the most common indices of functional diversity (Schleuter *et al*. 2010; Ricotta *et al*. 2016).

Consequently, the BNI shares a number of characteristics with Rao’s Q. Both indices are primarily based on pairwise distances between species, which are calculated from relevant attributes of species, such as functional trait values or phylogenetic distances. In the same way that pairwise distances are weighted by relative abundances in Rao’s Q, pairwise distances are weighted by a pairwise temporal coexistence coefficient in the BNI. This temporal coefficient is calculated based on the estimated residence time of each species in the reference area and captures how long pairs of species have coexisted in the area. For example, if a given pair of species consists of a native and a recently introduced species, their pairwise trait distance will be weighted more heavily than the distance between a native and another non-native which arrived earlier in the area. This temporal coefficient allows us to take into account the temporal erosion of novelty in a community, and differentiate between non-natives in such a way that a recently introduced species may be seen as “more novel” compared to the established non-native species.

We describe how to calculate the BNI from various data sources, and how it associates with traditional measures of biotic novelty, abiotic novelty, species richness and functional diversity. By presenting simulations and two case studies, we show that this new method to quantify biotic novelty is intuitive and versatile, as it is easily adaptable to datasets of different scale, scope and resolution. We demonstrate in this paper that the BNI framework is a helpful tool whenever the assessment of novel species assemblages or communities is needed, which may not only be useful in invasion ecology, but also in global change ecology, restoration ecology or urban ecology.

## Methods

### The new index of biotic novelty

There are seven steps to calculate the BNI: (1) obtaining a trait matrix, (2) converting the trait matrix into a distance matrix, (3) obtaining species’ first records, (4) converting the first records into a temporal coexistence matrix, (5) weighting the distance matrix by the temporal coexistence matrix, (6) multiplying the distance matrix by the species’ relative abundance (optional), and (7) calculating the sum of all pairwise comparisons from the distance matrix (Fig. 2). The resulting BNI is expressed as:

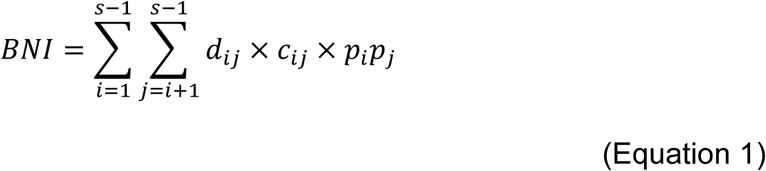

where *d*_*ij*_ is the distance between species *i* and *j, c*_*ij*_ is the temporal coexistence coefficient of species *i* and *j* in the local area, and *p*_*i*_*p*_*j*_ are the relative abundances of species *i* and *j*. Note that the equation of the BNI corresponds to the calculation of Rao’s Q (Rao 1982; Botta-Dukát 2005), but with the temporal coexistence coefficient *c*_*ij*_ added to the product term. The steps 1, 2, 6 and 7 are standard multivariate methods to obtain Rao’s Q; steps 3, 4 and 5 are the implementation of the temporal coexistence component. Both components are explained in detail in the following sections.

**Figure 2:**
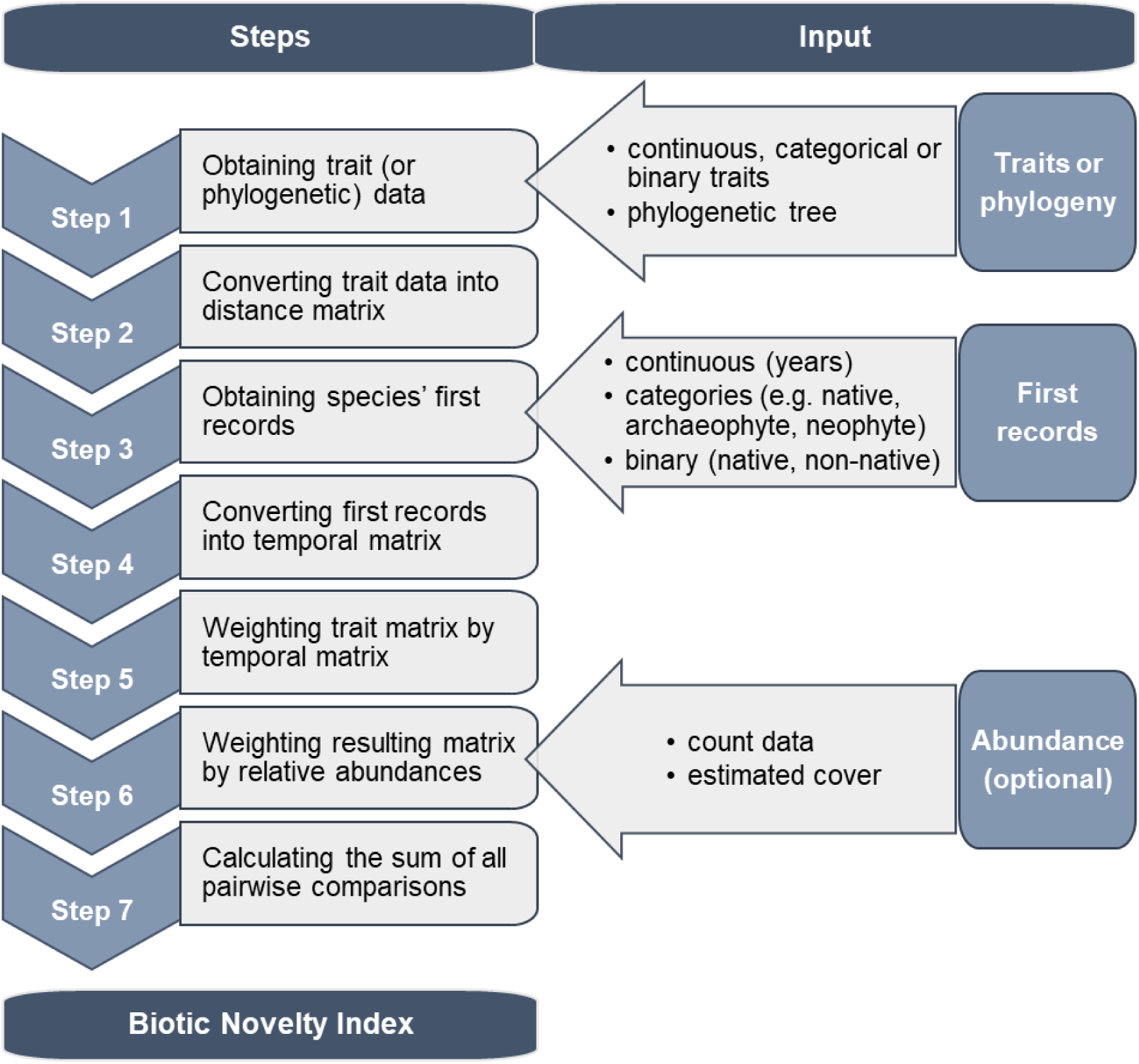
Standardized procedure for calculating the biotic novelty of a community with the Biotic Novelty Index (BNI).

### The functional diversity component

The general rule to calculate functional diversity indices is that traits must be linked to the function(s) of interest. For instance, specific leaf area, maximum growth rate and leaf nitrogen concentration are important components of plant functional diversity when primary production is the process of interest (Garnier *et al*. 2004; Wright *et al*. 2004). Similarly, the choice of traits for the BNI can be related to the novelty aspects of interest. For example, if the aim is to assess the biotic novelty of an invertebrate herbivore community, feeding preference, feeding type (e.g. chewing or sucking) and the number of generations per year are traits where novelty could play a relevant role for the consumed plant. If some traits are more important for evaluating biotic novelty than others, they should be given greater weights in the trait matrix. Careful decisions about which traits to include and how to weigh them depends on the purpose to which the index will be applied and should rely on expert knowledge of the system (Laliberté & Legendre 2010). Traits can be continuous (e.g. leaf nitrogen concentration), binary variables (e.g. legume or non-legume) or categorical (e.g. flower color).

Distance measures calculate the difference between pairs of species based on their characteristics (e.g. functional traits). There are many distance measures to choose from, but two are most commonly used on trait datasets: the Euclidean distance and the Gower distance (Laliberté & Legendre 2010). The Euclidean distance is calculated on complete and continuous trait datasets, and emphasizes absolute differences (Poos *et al*. 2009), while the Gower distance has the advantage that it allows incomplete data sets and mixed (categorical, ordinal, continuous) data types (Gower 1971; Laliberté & Legendre 2010).

### The temporal coexistence component

In the BNI, pairwise trait distances are weighted by a pairwise temporal coexistence coefficient. The first step in calculating this coefficient is to define whether each species belongs to the historical native species pool or not. Second, we use information such as first records (or time of establishment) of the non-native species in the local region. This information can be obtained either from publications (e.g. Seebens *et al*. 2017 collected first records of alien species worldwide: http://dataportal-senckenberg.de/database/metacat/bikf.10029/bikf), regional databases (e.g. the BiolFlor database for plants in Germany, Klotz *et al*. 2002), or expert knowledge. For native species, time of establishment needs to be estimated as well (e.g. for many plant species in Central Europe a reference to the end of the last glacial period will be reasonable). From this information, the residence time for each species is calculated. The residence time tells us how many years before today each species was introduced or had been established. For example, a species that was introduced in 1719 has a residence time of 300 years in the year 2019 (the current year). Next, resident times are normalized between the oldest residents (i.e. native species) and the newest arrivals, bringing them into the range [0,1] by the following calculation:

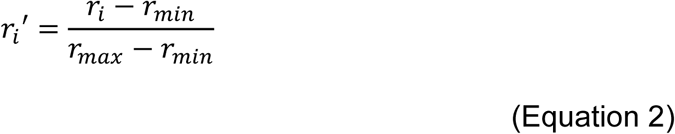

where 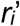 is the normalized residence time of species *i, r*_*i*_ is the residence time of species *i* (in years), *r*_*min*_ is the minimum residence time of all species (i.e. the newest arrival) and *r*_*max*_ the maximum residence time of all species (i.e. residence time of native species). Once the normalized residence time is calculated for each species, for each pair of species the temporal coexistence coefficient can be calculated as follows:

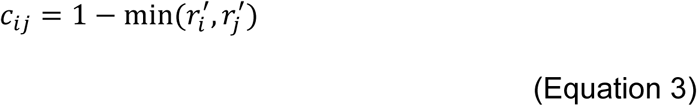

where *c*_*ij*_ is the temporal coexistence coefficient of species *i* and *j*, 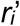 is the normalized residence time of species *i* and 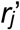 is the normalized residence time of species *j*.

Note that the minimum of both normalized residence times is used in equation 3 because the latest arrival in the species pair determines how long both species have coexisted in the focal area. For example, if the two species have residence times of 300 and 100 years, respectively, their temporal coexistence in the focal area is 100 years. We then take the complement of the minimum normalized residence time in equation 3, such that the coefficient is maximized when species have had the lowest local coexistence time (i.e. maximum novelty). Eventually, the temporal coexistence coefficient is calculated for each possible species pair and a new temporal matrix can be constructed with the same dimension as the trait distance matrix described before. The values of the temporal matrix range between 0 and 1 (due to the normalization step given in equation 2) and functions as weighting factor for the trait distance matrix. In this way, trait differences between species with low coexistence time are weighted heavily, whereas trait differences between species coexisting for millennia (such as a pair of native species) will be given no weight in the BNI.

### The BNI as a framework

The BNI is in essence the sum of two components: the mean functional distance between novel species in the community, and the mean functional distance between native and novel species. Furthermore, we can show that the BNI is an additive partition of Rao’s Q (see supplementary material S1 for details). According to this partitioning, we can express the BNI relative to Rao’s Q, and define a standardized version of the BNI as:

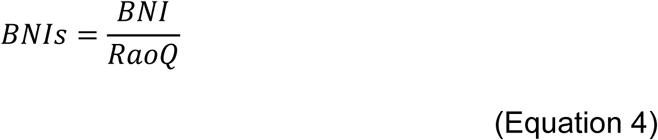

This standardized version is a proportion of Rao’s Q, which can be described as the proportion of functional diversity contributed by novel species pairs (for an application, see the simulations below and in supplementary material S2). A detailed R code, that helps the user to calculate the BNI and the BNIs, is provided in supplementary material 3.

We purposely refer to the BNI as a framework because it is built upon the idea to combine two relevant aspects into one measure, which can be easily adapted to the needs of the user (e.g. by adding or replacing relevant components) depending on the goal of the study. For example, the BNI as described above captures the functional novelty of communities because it uses functional traits to calculate differences between species. However, if the user aims to assess phylogenetic aspects of novelty, or to compare phylogenetic aspects with functional aspects, then the functional diversity component of the BNI may be replaced with a measurement of phylogenetic distances between species (see case study 2 for an application). While phylogeny has sometimes been used as a proxy for functional or ecological niche differences between species (Webb *et al*. 2002; Helmus *et al*. 2007; Cadotte *et al*. 2009), it has become clear that phylogenetic distances are, at best, an imperfect proxy (Emerson & Gillespie 2008; Mason & Pavoine 2013). Calculating the BNI using phylogenetic distances may be useful in cases when trait data are difficult to obtain or the evolutionary history and relatedness of species are the focus of interest (Gerhold *et al*. 2015).

While the temporal component of the BNI was designed to use species residence times as the most accurate way to weigh the novelty of species interactions, there will often be situations where dates of first records are imprecise, incomplete or even entirely missing. For these cases, we suggest the use of temporal categories to characterize each species in the community. The generation of these categories, for example, could be based on corresponding decades or centuries. Another approach would be to adopt already existing temporal categorizations such as the three-level classification scheme of European plant species by Schroeder (1968): non-native species are classified according to their time of human introduction, either before Europe’s discovery of the New World in 1492 (archaeophytes or more generally archaeobiota) or after 1492 (neophytes, neobiota). Species that colonized a given area after the end of the last glacial period without human assistance are classified as native (see case study 1 for an application). “Neonative” species could be added as another category for species establishing due to climate change in the Anthropocene, i.e. since the middle of the 20^th^ century (Essl *et al*. 2019). If even these data are not available, the user may opt for the most basic categorization method which classifies species as either native or non-native (i.e. a binary categorization). In this case, the corresponding temporal coexistence coefficient would be either 0 for pairs of native species, or 1 for pairs involving at least one non-native species.

The BNI as described above is a multispecies approach since it captures the functional novelty of communities and species assemblages. However, by modifying the BNI equation, it is also possible to focus on the biotic novelty of particular novel target species in relationship to the interacting resident species. A similar approach was proposed by Saul & Jeschke (2015), which consider the implications of different degrees of eco-evolutionary experience of interacting resident and novel species.

### Simulations

Simulations of plant communities were used to explore the behavior of the BNI in different scenarios of functional diversity and biological invasion. We randomly generated a regional pool of 250 species, with 70 % natives and 30 % non-natives. In order to spread the simulated residence times realistically, we followed the three-level classification of European plant species described before, and separated non-natives into long established non-natives (e.g. archaeophytes, 15 % of all simulated species) and recently arrived non-natives (e.g. neophytes, 15 % of all species). We attributed mean dates of arrival for each species based on these categories: 8518 years for natives, 2786 years for archaeophytes, and a uniformly random generated year of arrival since 1492 for the neophytes. The mean dates of arrival for natives and archaeophytes originate from the respective class limits of natives and archaeophytes in the Berlin/Brandenburg area, i.e. around 10,000 BC (end of the last glacial period) for natives and around 3,000 BC for the introduction of the first archaeophytes (Haas, Giesecke, & Karg, 2003). Next, we randomly generated functional trait values for each species. Three continuous traits were sampled from normal distributions, whose mean and variance were determined according to one of four non-native trait scenarios.

Since the BNI is designed to capture functional novelty, we explored scenarios where neophyte species are bringing different functional trait values from the historical resident pool of species (i.e. natives and archaeophytes pooled together). We present four trait scenarios: (1) traits for all species are sampled from the same distribution; (2) traits of neophytes have on average higher values than the residents (i.e. different mean); (3) traits of neophytes occupy a wider range of values than the other species (i.e. different variance parameter); and (4) traits of neophytes have both a different mean and a different variance than the traits of residents (cf. supplementary material 2 for additional scenarios). We then assembled 100 communities by sampling randomly from the simulated species pool. The number of species sampled per community was assigned randomly following a Poisson distribution (lambda = 25). In order to generate a gradient of increasing biological invasion for each simulation, the 100 communities were forced to integrate an increasing proportion of neophytes (0 %, 25 %, 50 %, 75 %, 100 %). Simulations were repeated 500 times, with incremental changes in parameters for each scenarios every 20 simulations (cf. supplementary material 2). We calculated Rao’s Q, the BNI and the BNIs for each simulated community. All simulations and calculations were done in R version 3.6.0 (R Core Team 2019), and all code is included in supplementary material 3.

### Case study 1: Biotic novelty of plant communities along an urbanization gradient

To illustrate the strengths and weaknesses of the newly proposed measure, we analyzed changes in biotic novelty along an urbanization gradient in dry grassland communities in Berlin, Germany. This vegetation type spans a range of near-natural to strongly human-shaped sites throughout the city. For this reason, urban dry grasslands have been selected as a model ecosystem within the CityScapeLabs, an experimental platform with a network of 56 permanent plots, established for the evaluation of biodiversity in urban environments. From April 18^th^ to May 19^th^ 2017, vegetation surveys were carried out in a 4 × 4 m plot within each of the 56 grasslands, recording the abundance (percent cover) of 234 vascular plant species. Trait data for the calculation of the BNI and Rao’s Q were extracted from the TRY database (Kattge *et al*. 2011) and the BiolFlor database (Klotz *et al*. 2002). We used data for twelve plant functional traits (plant height, specific leaf area, life form, flower color, flower class, clonal growth organs, length of dispersal unit, seed mass, leaf area, leaf nitrogen content, nitrogen fixation and mycorrhizal infection). Information on the first record of neophytes is based on the atlas of the Berlin flora (Seitz *et al*. 2012). All other species were classified as native or as archaeophytes (introduced by human agency before 1492) according to the BiolFlor database (Klotz *et al*. 2002). Note that exact first record information (e.g. dates) were only available for neophytes, but not for archaeophytes, nor native species, which is a typical situation of data availability for plant species in Europe. Hence, we used for these two categories a mid-range value for each species in the respective category and the exact first records for neophytes only. The mid-range value for natives and archaeophytes was calculated from the respective class limits in the focal area, i.e. around 10,000 BC (end of the last glacial period) for natives and around 3,000 BC for the introduction of the first archaeophytes in the Berlin/Brandenburg region (Haas *et al*. 2003). This resulted in an estimated residence time of 8518 years for natives and 2786 years for archaeophytes.

To analyze the relationship between the biotic novelty of plant communities and the level of urbanization (as a driver of ecological novelty), we applied a commonly used indicator of urbanization: the percentage of sealed surfaces (i.e. impervious soils) in the surrounding landscape (Lu & Weng 2006; Schwarz 2010). We calculated the mean percentage of sealed surfaces in a 500 m buffer area around each of the 56 plots using publicly available urban habitat maps from the Berlin Senate Department for Urban Development and Housing and QGIS 2.18.0 (QGIS Development Team 2016). Relationships of the BNI and the BNIs with the percentage of sealed surfaces, Rao’s Q and species richness were analyzed with linear models. All calculations were carried out using R version 3.4.3 (R Core Team 2017).

### Case study 2: Biotic novelty of co-occurring vascular plants in Germany

The second case study demonstrates the application of the BNI in conjunction with big datasets. Here, we aimed to calculate the BNI for co-occurring vascular plants in Germany and to evaluate how their biotic novelty is spatially related to the extent of urban areas. It is a feature of this case study that it extensively used freely accessible data from online databases. From the Global Biodiversity Information Facility (GBIF: The Global Biodiversity Information Facility 2019) we downloaded the occurrence dataset ‘Flora von Deutschland (Phanerogamen)’ which includes 9,577,887 records of 5,721 vascular plant species in Germany (Bundesamt für Naturschutz / Netzwerk Phytodiversität Deutschland 2018). These occurrence records are aggregated in 11 × 11 km grid cells of the grid of topographic maps (TK 25, scale 1:25000), which are officially used for the design of species distribution maps in Germany. We used phylogenetic pairwise distances to calculate the BNI. In this case, the BNI thus captures phylogenetic novelty rather than the functional novelty we calculated in our simulations and in case study 1. To do so, we pruned the extensive phylogeny ‘Daphne’ (Durka & Michalski 2012) for our species set. Daphne is a dated phylogeny of a large European flora for phylogenetically informed ecological analyses. Information whether a plant species is native or non-native in Germany plus information on first records for neophytes were obtained from the BiolFlor (Klotz *et al*. 2002) database. We calculated the BNI for each of the 3,003 grid cells and created a map using QGIS version 3.2.1 (QGIS Development Team 2018). A second layer, which indicates the extent of urban areas based on MODIS satellite data (Schneider *et al*. 2009) was added to the map. All calculations were carried out using R version 3.4.3 (R Core Team 2017) and the R package ‘picante’ (Kembel *et al*. 2010) for phylogenetic tree pruning.

## Results

### Simulations

Simulations showed that the BNI varies broadly with the proportion of non-native species and with the size of trait differences between species (Fig. 3). Overall, as long as neophytes made up less than 50 % of the relative abundance of species in the community, the BNI increased monotonously as more neophytes were added. Beyond this point, however, the BNI did not always increase with the proportion of neophytes. Its behavior depended on how much pairwise trait variance the neophytes were bringing to the community, relative to the resident species.

**Figure 3:**
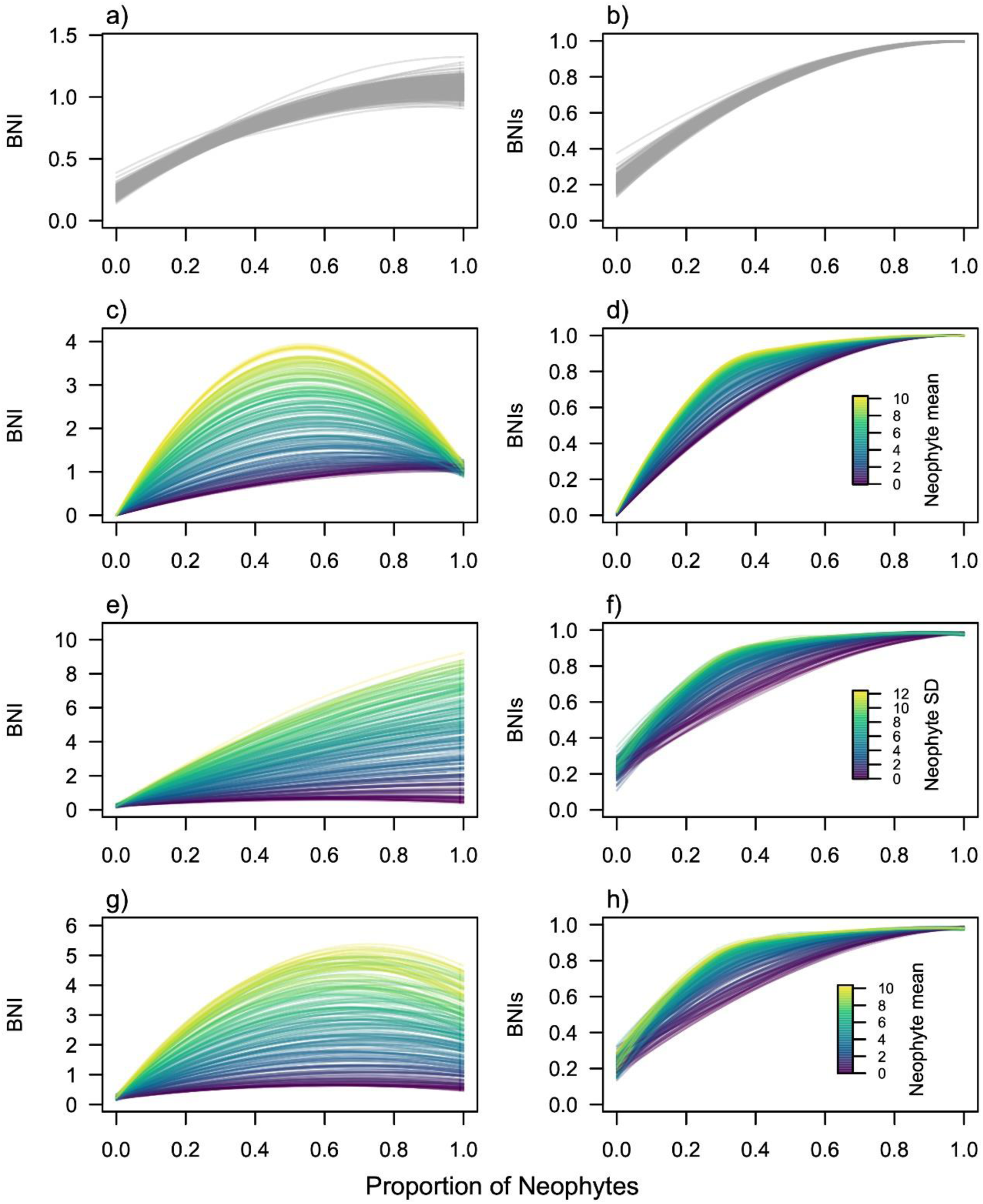
Variation of the Biotic Novelty Index (BNI) and its standardized value (BNIs) in four simulation scenarios. Communities were simulated with an increasing proportion of recently introduced non-native plants (neophytes). Scenarios explore different parameters (mean and SD) of the normal distribution from which species traits for neophytes were sampled. In the first scenario (a, b), traits of native and non-native species follow the same normal distribution (trait mean = 0, SD = 1). In scenario 2 (c, d), the mean trait values of neophytes are increasingly different from the natives (colors represent variation in neophyte trait mean from 0 to 10; SD = 1). In the third scenario (e, f), natives and neophytes have the same trait mean (mean = 0), but neophyte trait SD increases from 0 to 10. In the fourth scenario (g, h), both the mean and SD of neophyte trait distributions increase together from 0 to 10 and 0 to 5, respectively. Lines represent LOESS regressions fitted on the 100 simulated points corresponding to one simulation run.

In scenario 1, when neophytes were not on average functionally different from natives, the BNI increased monotonously with the proportion of neophytes (Fig. 3a). This is because, in this scenario, the mean pairwise trait differences (i.e. Rao’s Q) remained constant, while the contribution of neophytes increased with their relative abundance in the community. The BNI simulation curve tended to saturate at high neophyte proportions as new neophyte species were less likely to add new trait differences.

In scenario 2 and 4, when neophytes were on average functionally different from the residents, the simulated BNI often showed a humped-shaped curve, with a maximum at intermediate proportions of neophytes (Fig. 3c, g). This pattern is due to the fact that the BNI is based on mean pairwise differences between species, which reaches its maximum when one half (i.e. the neophytes) of the community is different from the other (i.e. the resident species). A similar pattern could be observed for Rao’s Q (details provided in supplementary material 2). Beyond this mid-point, the amount of trait variance among the neophytes (SD_neo_) determined the behavior of the BNI. As illustrated in scenario 3 and 4 (Fig 3e, g), as long as the trait values of the neophytes were more variable than those of the resident species (SD_neo_ > SD_residents_, with SD_residents_ = 1), the BNI increased monotonously with the proportion of neophytes and the amount of variance in neophyte traits. On the other hand, if neophytes had a lower trait variance (i.e. they were more similar amongst themselves) than the residents (SD_neo_ < SD_residents_), then the BNI tended to decrease with the proportion of neophytes.

These simulations illustrate how the BNI captures the absolute contribution of novel species to functional diversity. As a consequence, communities composed of functionally very similar non-natives will tend to have low functional diversity and a low BNI. Interpretation of BNI values must therefore consider the relative abundance (or proportion) of the non-native values in the community.

By contrast, the standardized value of the BNI (BNIs) showed no such changes in behavior across scenarios. The BNIs increased monotonously with the proportion of neophytes. The rate of increase was always higher than 1, with steeper curves generated by neophyte traits being different on average from residents (scenarios 2 and 4), or with higher variance than residents (scenarios 3 and 4).

### Case study 1: Biotic novelty of plant communities along an urbanization gradient

The observed BNI values for the 56 Berlin grassland plots ranged from 0.002 to 0.092 and had a mean at 0.020 ± 0.016 SD. The plot with the lowest BNI value contained 13 species of which 12 were native and 1 was non-native, specifically an archaeophyte species. The plot with the highest BNI value contained 32 species of which 19, 6 and 7 were native, archaeophytes and neophytes, respectively. Statistical analyses of the BNI across the 56 plots indicated that the BNI was positively related to the urbanity indicator sealed surface area (Fig. 4). 15% of the variation in the BNI was explained by the percentage of sealed surfaces around the plots (*P* = 0.003, Fig. 4a). However, there were no significant relationships detectable between the sealed surface area and traditional measures of biotic novelty, i.e. the number of nonnative species (*R*^*2*^ = 0.01, *P* = 0.443, Fig. 4b) or their proportion (*R*^*2*^ = 0.04, *P* = 0.130, data not shown). Further, when considering total functional diversity (expressed as Rao’s Q), we also identified a positive relationship with the sealed surface area (*R*^*2*^ = 0.08, *P* = 0.040, Fig. 4c), but less strong than the one for the BNI. Finally, we investigated how the BNI varies independently of the variation in Rao’s Q by calculating the standardized version of the BNI. The standardized BNI (BNIs) showed a similar relationship with the sealed surface area (*R*^2^ = 0.14, *P* = 0.004, Fig. 4d) than the non-standardized BNI.

**Figure 4:**
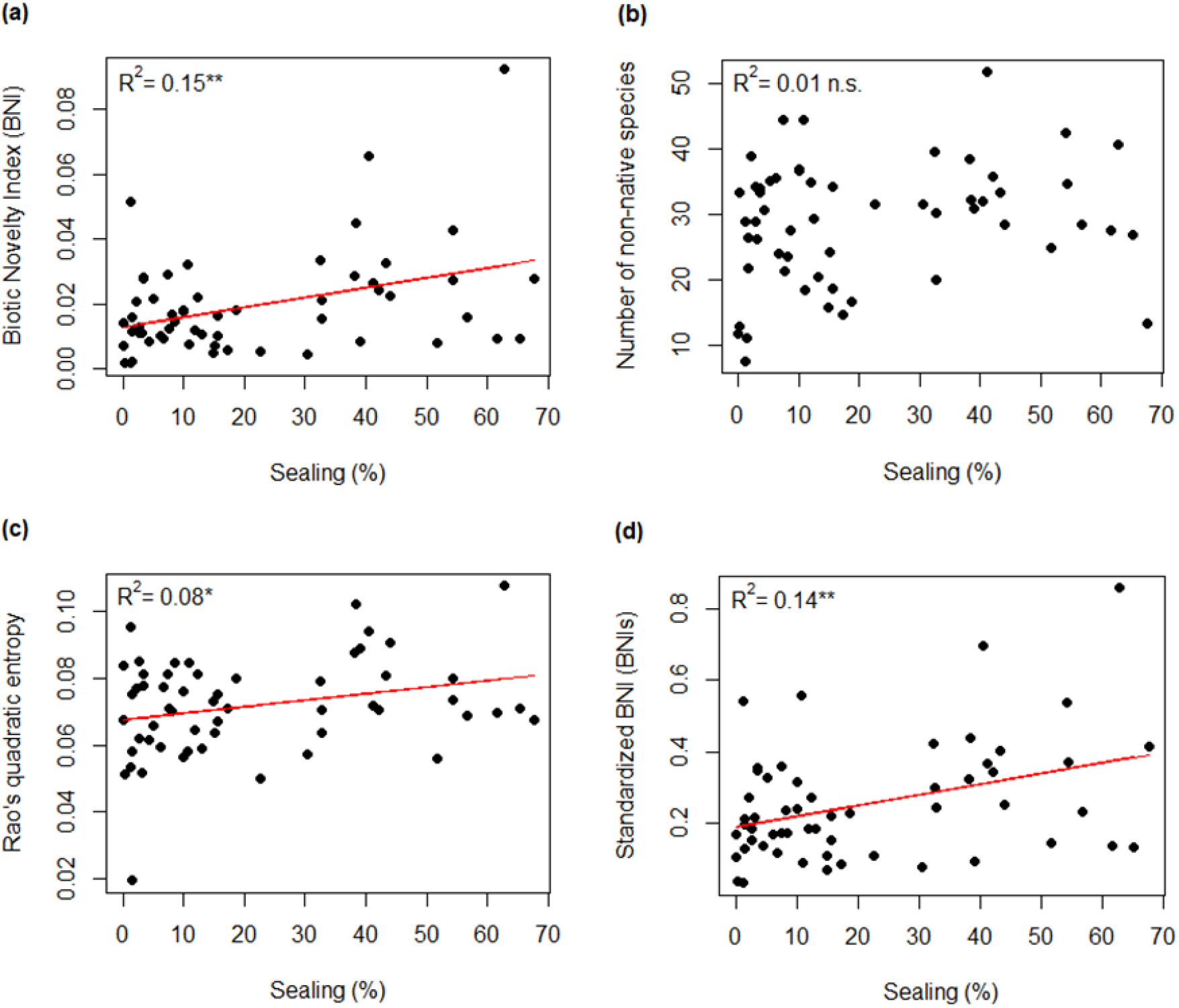
Case study 1 – relationships between the percentage of sealed surface area in a 500 m buffer zone around the 56 urban grassland plots and (a) the BNI, (b) the number of non-native species, (c) Rao’s Q as a measure of functional diversity, and (d) the standardized BNI. Asterisks indicate statistical significance using linear models (‘***’ = *P* < 0.001, ‘**’ = *P* < 0.01, ‘*’ = *P* < 0.05, ‘n.s.’ = *P* ≥ 0.05).

We were also interested in how the BNI associates with community parameters such as species richness and functional diversity. The BNI was not related to the total number of species in the plots (*R*^*2*^ = 0.05, *P* = 0.103, Fig. 5a), but showed a moderately positive relationship with the number of non-native species (*R*^*2*^ = 0.23, *P* < 0.001, Fig. 5b). On the other hand, the BNI was strongly positively related with the functional diversity (expressed as Rao’s Q) of all species (*R*^*2*^ = 0.43, *P* < 0.001, Fig. 5c), but weakly positively related to the functional diversity of the group of non-native species (*R*^*2*^ = 0.09, *P* = 0.028, Fig. 5d). The standardized version of the BNI (BNIs) showed almost identical relationships to all four community parameters (Fig. S4.1 in supplementary material S4).

**Figure 5:**
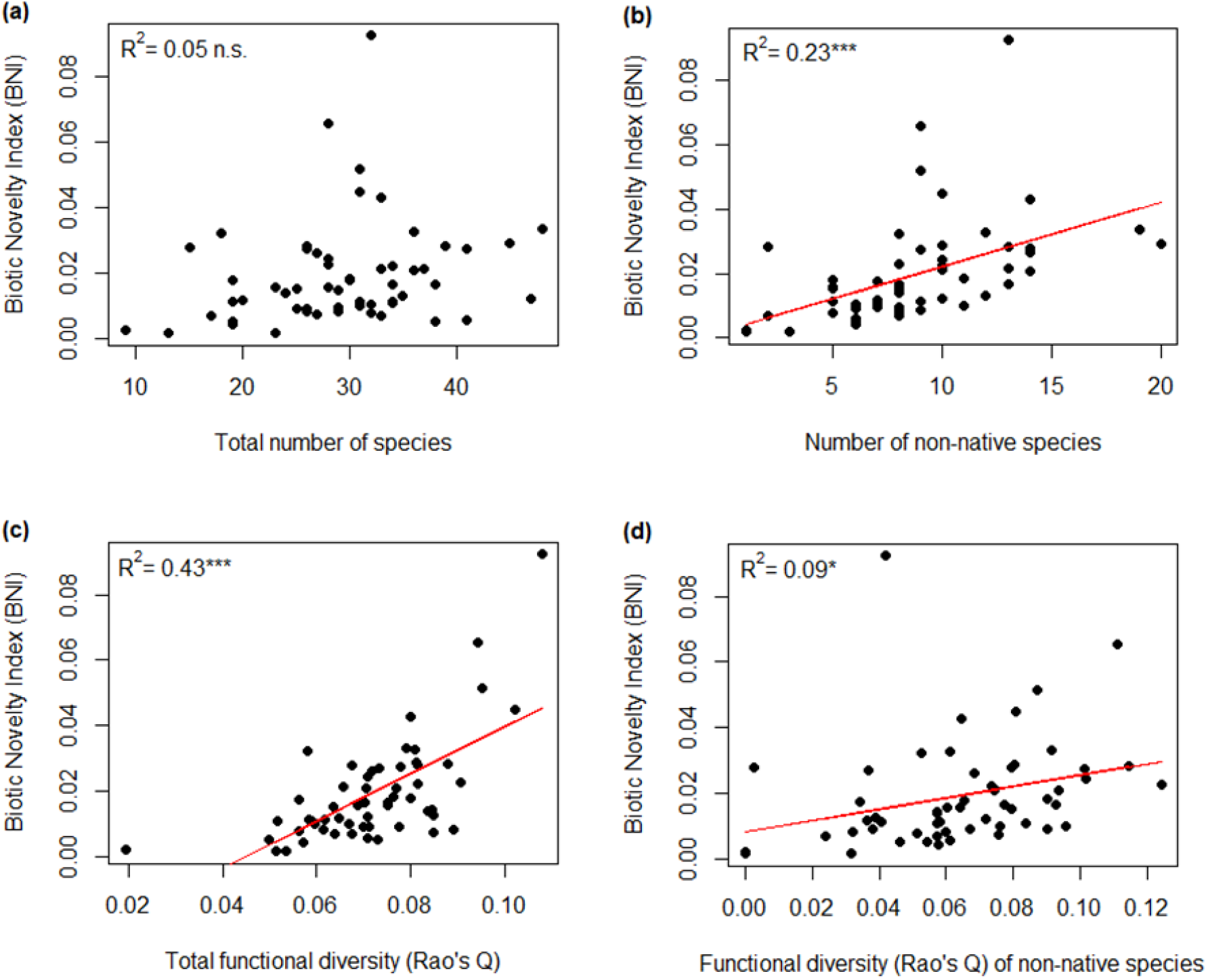
Case study 1 – relationships between the BNI and (a) the total number of species, (b) the number of non-native species, (c) Rao’s Q as a measure of functional diversity, and (d) the functional diversity of non-native species in the 56 urban grassland plots. Asterisks indicate statistical significance using linear models (‘***’ = *P* < 0.001, ‘**’ = *P* < 0.01, ‘*’ = *P* < 0.05, ‘n.s.’ = *P* ≥ 0.05).

### Case study 2: Biotic novelty of co-occurring vascular plant species in Germany

The nationwide assessment of biotic phylogenetic novelty identified large areas with high novelty in Germany, indicated by the distribution map and the slightly right-skewed histogram of the BNI (Fig. 6). The BNI values ranged from 0 (at Zugspitze, the highest mountain in Germany) to 64.18 (in Leipzig, the most populous city in the German federal state of Saxony). Areas of very high novelty were clearly concentrated in and around urban areas: in addition to Leipzig, other areas of high novelty were the cities Cologne (62.72), Bamberg (62.39) and Mülheim an der Ruhr (62.15). The capital and largest city of Germany, Berlin, had the 9^th^ highest BNI (61.06). That the city surroundings also showed a higher extent in biotic novelty may be indicative for a spatial spillover effect from cities to adjacent areas. However, this effect seemed to be less pronounced in southern Germany. Areas of low novelty were visible predominantly in southern and partly in central Germany, presumably due to the ranges of the Alps and the central uplands, respectively, in these regions. The standardized version of the BNI (BNIs) showed an almost identical distribution map (Fig. S4.2 in supplementary material S4).

**Figure 6:**
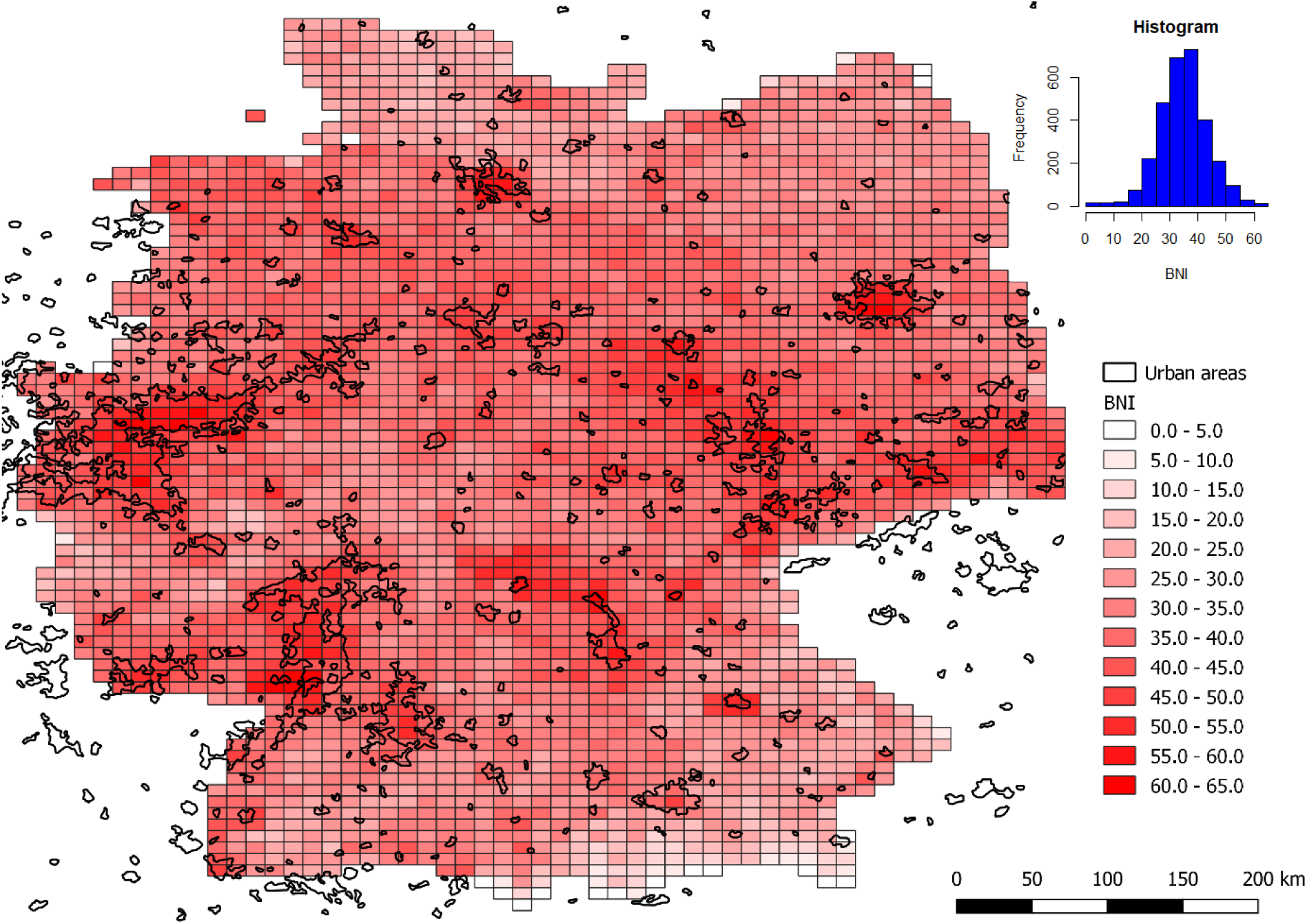
Case study 2 – biotic novelty of co-occurring vascular plants in Germany aggregated in 11 × 11 km grid cells calculated with the BNI. Areas outlined in black indicate the extent of urban areas based on MODIS satellite data (Schneider *et al*. 2009).

## Discussion

This study introduced the Biotic Novelty Index (BNI) and demonstrated its applicability as a framework to measure the ecological novelty of communities at different spatial scales. We regard ecological novelty as a continuous gradient ranging from historic or analog to novel (Heger *et al*. 2019) rather than a binary classification. Accordingly, we have designed the BNI to be able to gradually measure ecological novelty. More specifically, the BNI focuses on the biotic rather than abiotic component of ecological novelty (i.e. biotic novelty). It measures the extent of trait differences among novel and non-novel species and, simultaneously, takes temporal dynamics into account. Arithmetically, the BNI represents the expected functional novelty between two randomly picked individuals in the community. Further, we refer to the BNI as a framework because it is built upon the idea of combining two relevant aspects of a research field into one formula, which can be easily adapted to the needs of the user (e.g. by adding or replacing relevant components).

### The BNI captures novelty in both functional diversity and introduction history

We designed the BNI to combine two aspects of ecological novelty: historical novelty, captured by the sequence of arrivals of new species in a given region, and functional novelty contributed by the new species (Heger *et al*. 2019). Simulations show that the BNI does capture the latter aspect in a predictable manner: for a given proportion of non-native species, increasing trait differences between species increases the functional novelty of the community, and the BNI increases accordingly. However, the behavior of the BNI is not always linear in response to the first aspect, i.e. the proportion of non-native species. The BNI may be maximized at intermediate proportions of non-native species, when the most functionally different pairs of species (in our case the resident species vs. the neobiota) are also the most heavily weighted in the calculation, both by their relative abundances and by the temporal coefficient. This behavior of the BNI demonstrates its similarities with Rao’s quadratic entropy as a diversity measure. The following applies to Rao’s Q: when a new species is added to a given community and this species is functionally very similar or identical to the resident species, the addition of this new species results in a lower functional diversity of the community. Rao’s Q is thus maximized when the most different species in the community are in high abundance. To the BNI, this property translates in the following manner: when a non-native species is added to a given community and this species is very similar or identical to the other pre-existing non-natives in the considered traits, the addition of this non-native species may result in a lower BNI for the community. This behavior might be counterintuitive depending on the goal of the study and the user’s viewpoint on biotic novelty, which is why we also recommend calculating the standardized BNI values (BNIs).

The BNIs offers an additional description of biotic functional novelty of the community by quantifying the proportion of functional diversity (measured as Rao’s Q) that is contributed by novel species interactions in the community. The advantage of this standardization is that, by construction, it is monotonous with regard to increasing proportions of non-native species, and the size of trait differences. This standardized version may therefore provide a more objective measure to compare the level of biotic novelty between communities with different levels of functional diversity, or assembled from a different species pool. Nevertheless, the untransformed value of the BNI remains a valuable measurement when the goal is to quantify the absolute amounts of functional diversity contributed by novel species in a community. Depending on a study’s goal, we would recommend to use either of the two or both versions of the index in combination; the latter gives a fuller picture of variation in novelty across communities.

### Case studies

Both case studies revealed strong connections of biotic novelty, as measured with the BNI or BNIs, with abiotic novelty. The first study showed that the BNI of 56 dry grassland plant communities in Berlin was positively related to the observed urbanity indicator (i.e. percentage of sealed surfaces). This is not surprising, as previous studies demonstrated that the construction and expansion of towns and cities promote the loss of native species and their replacement by non-native species (Chocholoušková & Pyšek 2003; Standley 2003; DeCandido *et al*. 2004; Tait *et al*. 2005; Knapp *et al*. 2010). Further, spatial analyses often show that, for many taxa, increasing intensity of urban activity causes non-native species to increase in abundance and species richness while native species decline (McKinney 2001, 2006; Godefroid & Koedam 2007; Kowarik 2008). For example, in rural floras around Berlin, there are less than 20 % non-native plant species, but from the outskirts to the city center of Berlin, the percentage of non-native species increases from about 30 to 50 % of all species (Kowarik 2008). The high non-native species richness of urban floras has often been explained by increasing importation of non-native individuals and favorable habitat for the establishment of non-native species (McKinney 2006). However, in the present study system, this relationship between increasing urbanity and non-native species richness was not supported since we found no relationship between the sealed surface area and non-native species richness (nor their proportion on total species richness). This finding underlines that the BNI captures different aspects of biotic novelty than the plain number and/or proportion of non-native species.

Our analyses also showed a strong relationship of the BNI with Rao’s Q. This was expected, given that the BNI is actually an additive partition of Rao’s Q (see supplementary material 1 for details). Several recent studies also examined whether invasions of non-native species change the structure of native communities by increasing or decreasing functional diversity (Castro-Díez *et al*. 2016; Loiola *et al*. 2018; de la Riva *et al*. 2019). These measures that compare invaded and uninvaded communities functionally and calculate the magnitude of change share a similar basis with the BNI. However, the BNI includes all possible species pairings weighted by the temporal coexistence coefficient rather than a comparison of categories (which Castro-Díez *et al*. 2016; Loiola *et al*. 2018 and de la Riva *et al*. 2019 do). These conceptual differences in how biotic novelty is assessed were reflected in the result that the BNI was only weakly positively related to the functional diversity of the group of nonnative species (Fig. 5d).

Further, by applying the standardization of the BNI (the BNI in proportion to Rao’s Q), we showed in the first case study that the BNI was not driven by the inherent variation in functional diversity along the urbanity gradient (since BNI and BNIs varied to a very similar extent along the gradient). As shown in our methods section, this standardization of the BNI can be easily applied by the user for a validation of the BNI results.

The second case study demonstrated the applicability of the BNI to nationwide datasets. The grid-cell map showed that areas of very high novelty of vascular plant species were predominantly concentrated in and around urban areas in Germany, which is partially in line with former nationwide assessments of vascular plants in Germany (Kühn *et al*. 2004) and the UK (Botham *et al*. 2009). These studies described that neophytes were very strongly associated with urban land cover, but do not appear to be spreading out of urban habitats into the wider countryside. Our finding that the BNI is also higher around urban areas might be due to spread of novel species along transportation pathways, such as roads (von der Lippe & Kowarik 2008) and rivers (Maskell *et al*. 2006), which connect cities and are located in corresponding grid cells in the map.

We observed on the grid-cell map that areas of low novelty were visible predominantly in southern Germany and partly in central Germany, which coincidences with mountain ranges in Germany. Previous studies also showed that non-native species richness typically declines along elevational gradients (Alexander *et al*. 2011; Seipel *et al*. 2012; Averett *et al*. 2016). This pattern has been explained by two factors: (1) special adaptations are required to invade extreme environments (Alpert *et al*. 2000; Pauchard *et al*. 2009; Alexander *et al*. 2011), making mountains inherently resistant to invasions; and (2) anthropogenic disturbance decreases with increasing elevation, leading to fewer species introductions (i.e. lower propagule pressure) and also higher resistance to invasions (Arévalo *et al*. 2005; Averett *et al*. 2016).

We are aware that analyzing a dataset with the extent of our second case study is not free of concerns. For example, the large grid-cell size (11 x 11 km) and the spatial autocorrelation of grid cells (Kühn *et al*. 2004) may be problematic sources of error. Sampling bias (i.e. there are more botanical institutes and experts in urbanized areas than in less urbanized areas) and other potential explanatory variables (e.g. geological types of grid cells) may play important roles for such an analysis as well. However, since it is the scope of this paper to demonstrate possible applications of the BNI rather than disentangling various factors that structure biotic novelty, we refrained to perform complex statistical analysis and chose to present a map without underlying models. Therefore, it is up to future studies to focus on this demanding task.

## Conclusions

Human-induced changes are generating novel communities composed of new combinations of species which may result in increased biotic novelty. Previous methods for quantifying biotic novelty, such as counting the number of non-native species, appear limited in that they do not consider whether these new species are functionally novel, or how long these species have been residents, possibly over- or underestimating the amount of novelty contributed by these new species. Our framework of measuring biotic novelty may have an advantage over a number of measures by combining these relevant aspects of biotic novelty into a single formula, accompanied by a straightforward standardization method. It allows for a nuanced comparison of communities, as it considers the trait differences between species. It is also versatile, since it allows species differences, hence novelty, to be measured in different ways according to the focus of the study. It is a helpful tool whenever the assessment of novel species assemblages is needed, which is not only the case in invasion ecology, but also in global change ecology, restoration ecology or urban ecology. We encourage further use and development of the BNI framework for different purposes in the future.

## Supporting information

Supplementary material 1 - Mathematics

Supplementary material 2 - Simulations

## Acknowledgements

This work was funded by the German Federal Ministry of Education and Research BMBF within the Collaborative Project “Bridging in Biodiversity Science BIBS” (funding number 01LC1501A-H). JMJ was additionally supported by the Deutsche Forschungsgemeinschaft (DFG; JE 288/9-2). We thank Anne Hiller for providing data on the characteristics of the urban matrix surrounding the study sites. We thank Gabriela Onandia, Arthur Gessler, Johannes Müller, Mark-Oliver Rödel, Stephanie Niemeier, Silvia Keinath, Hans-Peter Grossart and Silvia Eckert for valuable discussions during meetings of the Work Package 5 in the BIBS project.

