## Supplementary material 1 - Mathematics for "A multidimensional framework for measuring biotic novelty: How novel is a community?"

### Supplementary material S1 – The Biotic Novelty Index as a partition of Rao's quadratic entropy

#### 1. BNI calculation

As a reminder, here is the formula for the Biotic Novelty Index (BNI) as presented in the main text of the article:

$$BNI = \sum_{i=1}^{s-1} \sum_{j=i+1}^{s-1} d_{ij} \times c_{ij} \times p_i p_j \qquad \text{(Equation 1)}$$

where $d_{ij}$ is the distance between species *i* and *j*, $c_{ij}$ is the temporal coexistence coefficient of species *i* and *j* in the local area, and $p_i$ and $p_j$ are the relative abundances of species *i* and *j* in a given species assemblage.

The temporal coexistence coefficient is calculated based on the normalized residence time of species:

$$r_i' = \frac{r_i - r_{min}}{r_{max} - r_{min}} \qquad \text{(Equation 2)}$$

where $r_i'$ is the normalized residence time of species *i*, $r_i$ is the residence time of species *i*, $r_{min}$ is the minimum residence time of all species and $r_{max}$ the maximum residence time of all species. Once the normalized residence time is calculated for each species, for each pair of species the temporal coexistence coefficient can be calculated as follows:

$$c_{ij} = 1 - \min(r_i', r_j') \qquad \text{(Equation 3)}$$

where $c_{ij}$ is the temporal coexistence coefficient of species *i* and *j*, $r_i'$ is the normalized residence time of species *i* and $r_j'$ is the normalized residence time of species *j.*

**2. BNI as a partition of Rao's Quadratic Entropy**

The BNI index can be expressed as an additive partition of Rao's quadratic entropy (Rao's Q; Rao 1982):

$$RaoQ = \sum_{i=1}^{s-1} \sum_{j=i+1}^{s-1} d_{ij} \times p_i p_j \qquad \text{(Equation 4)}$$

$$= \sum_{i=1}^{s-1} \sum_{j=i+1}^{s-1} d_{ij} \times ( c_{ij} + 1 - c_{ij}) \times p_i p_j$$

$$= \sum_{i=1}^{s-1} \sum_{j=i+1}^{s-1} \left[d_{ij} \times c_{ij} \times p_i p_j + d_{ij} \times \left(1 - c_{ij}\right) \times p_i p_j\right]$$

$$= \sum_{i=1}^{s-1} \sum_{j=i+1}^{s-1} d_{ij} \times c_{ij} \times p_i p_j + \sum_{i=1}^{s-1} \sum_{j=i+1}^{s-1} d_{ij} \times \left(1 - c_{ij}\right) \times p_i p_j$$

$$= BNI + \sum_{i=1}^{s-1} \sum_{j=i+1}^{s-1} d_{ij} \times \left(1 - c_{ij}\right) \times p_i p_j \qquad \text{(Equation 4a)}$$

This partitioning of Rao's Q can be expressed as the sum of two components: the "novelty component" (BNI), and what we might call a "biotic familiarity component" (BFC), which corresponds to the amount of functional differences in the community with which species have coexisted for a long time:

$$RaoQ = BNI + BFC \qquad \text{(Equation 4b)}$$

**3. Simplified version of the partition**

This partitioning of Rao's Q into the BNI becomes easier to picture when considering a simplified case in which we only use information about the native vs. alien status of species, ignoring differences in residence time. In this case, the temporal coexistence coefficient is reduced to:

$c_{ij} = 0$ if *i* and *j* are both native

$c_{ij} = 1$ if *i* and/or *j* are alien

Therefore, $(1 - c_{ij})$ in **Equation 4a** would only be non-zero for cases where both species in the pair are native, which means the equation simplifies to:

$$RaoQ = BNI + RaoQ_{Natives} \quad \text{(Equation 4c)}$$

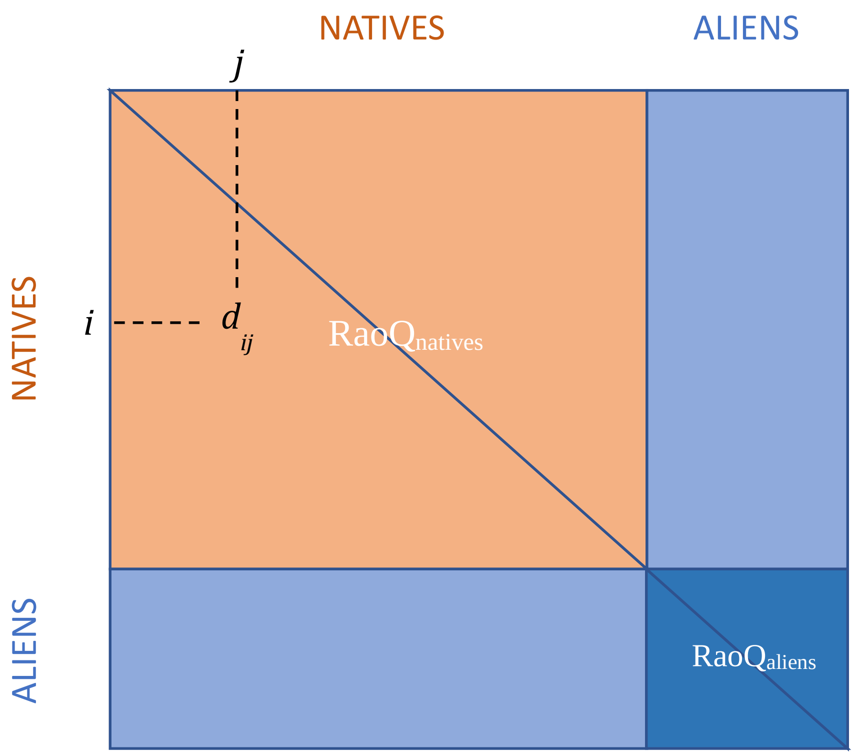


**Figure S1.1. Illustration of the additive partitioning of pairwise distances between native and alien species in a community, relevant to the simplified version of the BNI.** A matrix of pairwise species distances ($d_{ij}$; e.g. functional distance) is schematically repesented, with distances between native species represented in orange, and distances between alien species and the rest of the community in blue. In the simplified case of **Equation 4c**, BNI coresponds to the abundance-weigthed mean of all the blue distances, while $RaoQ_{Natives}$ is the weighted mean of all the orange distances. In **Equation 4d**, BNI is further partioned into $RaoQ_{aliens}$ (dark blue square) and the mean distance between natives and aliens (light blue areas).

In this simplified case, the “biotic familiarity component” (**Equation 4b**) is equal to the functional diversity (calculated as Rao’s Q) of native species in the community (Figure 1). Given the simplified values of $c_{ij}$ in this scenario, we can further decompose the BNI:

$$RaoQ = RaoQ_{Aliens} + 2 \times \sum_{i=alien} \sum_{j=native} d_{ij} \times p_i p_j + RaoQ_{Natives} \qquad \text{(Equation 4d)}$$

The middle component in **Equation 4d** corresponds to the mean of all pairwise distances between alien and native species in the community (light blue areas in **Figure 1**).

The BNI is in essence the sum of these two components: the mean functional distance between non-native species in the community (i.e. Rao’s Q for aliens only), and the mean functional distance between natives and non-natives. Taking into account the different residence times of alien species blurs this demarcation between natives and aliens, and provides additional nuance to the degree of novelty that a species contributes to the species assemblage.

**4. A standardized version of the BNI**

This additive partitioning of Rao’s quadratic entropy implies that we can calculate a strandardized version of the BNI (BNIs) with respect to Rao’s Q. This standardized index captures the proportion of functional (or phylogenetic) diversity contributed by novel pairwise interactions in a given species assemblage:

$$BNIs = \frac{BNI}{RaoQ} \qquad \text{(Equation 5)}$$

An assemblage made up entirely of natives will have a BNIs (and also a BNI) equal to zero, i.e. none of the funcitonal diversity is contributed by novel species. An assemblage made up entirely of species which arrived last year in the region will have a BNIs equal to 1, as the totality of the functional diversity in the assemblage is made up of novel species and equally weighted by $c_{ij}$ = 1.
