## Supplementary material 2 - Simulations for "A multidimensional framework for measuring biotic novelty: How novel is a community?"

We ran simulations to explore the behavior of the Biotic Novelty Index (BNI) and the standardized version of the BNI (BNIs). In each simulation, we generated a random regional pool of species and assembled 100 local communities distributed along a gradient of increasing proportion of neobiota. We explored different scenarios of trait distributions in the species pool with respect to species biogeographic origin (neobiota vs. older residents), and observed how the BNI and BNIs varied along the gradient of proportion of neobiotia in each scenario.

### 1. Community assembly simulation procedure

The simulation procedure performed the following steps:

1. Create a pool of 250 species.
2. Assign introduction status to each species: 75 % native, 15 % archaeobiota, 15 % neobiota.
3. Assign residence times for each species depending on the introduction status: 8518 years for all natives, 2768 years for all archaeobiota, and an integer randomly drawn between 1 and 526 years for each neobiota.
4. Assign random trait values to each species in the pool, following one of the trait scenarios (*cf. next section*)
5. Assemble 100 local communities, distributed along a gradient of proportion of neobiota. This gradient comprises 20 communities from each of the following proportions of neobiota: 0 %, 25 %, 50 %, 75 % and 100 %.

a. Assign species richness for each community. Values are sampled randomly from a Poisson distribution of parameter lambda = 25 (i.e. expected species richness was 25 species across all communities).
b. Draw randomly species from the species pool, in order to obtain the assigned species richness and proportion of neobiota.

6. Calculate the BNI, Rao's Q and the BNIs

This simulation procedure was repeated 200 times per scenario. Note that to simplify assumptions, species abundances are all considered to be equal in these simulated communities.

**2. Trait scenarios tested**

Much research has been done concerning the traits of non-native species which have been successful in establishing or becoming invasive in their new range. In some cases, non-native species exhibit different traits from the natives (e.g. Ordonez, Wright, & Olff 2010; Ordonez 2014), while in others they appear rather to blend in (e.g. Cross, Green, & Morgan 2015). Such patterns of trait differences between natives and introduced species are expected to vary according to traits, habitats and spatial scale (Hulme & Bernard-Verdier 2018).

We simulated different patterns of trait difference between native and non-native species in the simulated species pool from which all local communities were assembled. While we explored trait scenarios for the species pool, local community assembly was kept neutral with respect to traits.

At each simulation run, three different traits (T1, T2, T3) were randomly drawn from normal distributions and assigned to the 250 species of the species pool. Two of these traits were kept the same for all species, being sampled from the same centered normal distribution (mean = 0 and SD = 1). The third trait (T3) was sampled from a normal distribution whose parameters varied according to the origin of the species. For natives and archaeobiota (i.e. the “resident species”), T3 was always sampled from a centered normal distribution (mean = 0 and SD = 1). For neobiota, the third trait was sampled according to 4 types of scenarios (cf. Table S2.1):

1- No difference between natives and neobiota trait values.
2- Traits of neobiota are different on average from those of natives.
3- Traits of neobiota have a different variance than natives:
    a. Neobiota are generally more variable than natives.
    b. Natives are already very variable, and neobiota have a lower range than natives.
4- Traits of neobiota are different in average and in variance from natives. Two options were explored for this scenario:
    a. Mean and variance of neobiota traits covary positively (e.g. neobiota show less trait filtering than natives).
    b. Mean and variance of neobiota covary negatively (e.g. neobiota are highly filtered on this trait).

Each run of a scenario generates one species pool and 100 local communities. We ran the same scenario 200 times, generating 2000 communities. We calculated the BNI, Rao’s Q and BNIs (BNI/Rao’s Q) for each community at each run. The exact parameters of each scenario are indicated in **Table S2.1**.

**Table S2.1. Parameters of the normal distributions from which neobiota and native traits were drawn in each simulation scenario.** When ranges are indicated, it means that scenario simulations were repeated for 20 levels uniformly distributed along the range.

| Scenario | Description | Neobiota | | Natives + Archaeobiota | |
|---|---|---|---|---|---|
| | | *Mean* | *SD* | *Mean* | *SD* |
| **Scenario 1** | No difference | 0 | 1 | 0 | 1 |
| **Scenario 2** | Different mean neobiota traits | 0—10 | 1 | 0 | 1 |
| **Scenario 3a** | Different SD of neobiota traits | 0 | 0—10 | 0 | 1 |
| **3b** | Different SD of neobiota traits, natives already have a large trait SD | 0 | 0—10 | 0 | 5 |
| **Scenario 4a** | Increasing neobiota trait mean and SD | 0—10 | 0—5 | 0 | 1 |
| **4b** | Opposite variation of neobiota trait mean and SD | 0—10 | 5—0 | 0 | 1 |

### 3. Simulation results

Results from each simulation scenario are presented below in **Figures S2.1—S2.6**. For each scenario we present: (1) an example of the simulated trait values across native and non-native species in the most extreme simulated species pool (= yellow curve of BNI); (2) variation of the BNI along an increasing proportion of neobiota in the community; (3) the corresponding variation in Rao's Q; (4) the corresponding variation in the BNIs.

Across scenarios, we observe that the BNI generally tends to increase with the proportion of neobiota in the community (x-axis), and with the amount of functional difference added by neobiota compared to the natives (i.e. curves increasing from violet to yellow). However, in some cases the BNI follows a hump-shaped curve, with the BNI decreasing after reaching a peak at intermediate proportion of neobiota. The monotonous vs. quadratic shape of the BNI depends on the amount of variation added by neobiota to the community.

If neobiota traits are not different from native traits (Scenario 1; **Fig S2.1**), adding neobiota to the community does not increase functional diversity (i.e. Rao's Q remains constant) but the BNI tends to increase slowly reaching a maximum roughly equal to the standard deviation of traits in neobiota.

If neobiota traits are very different on average from the traits of native species, but show equal standard deviations (Scenario 2; **Fig. S2.2**) then the maximum BNI (and maximum Rao's Q) will be reached close to 50 % of neobiota in the community.

If neobiota are not different on average from natives, then differences in standard variation of traits among the neobiota will determine the shape of the relationship. In Scenario 3a (**Fig. S2.3**), and especially in Scenario 3b (**Fig. S2.4**), we observe that the BNI follows a hump-shaped curve as long as the standard deviation of neophyte traits is lower than that of the natives (maximum marked by red dots; $SD_{neo} < SD_{nat}$), but increases monotonously as soon as it becomes larger (maximum marked by black dots; $SD_{neo} \geq SD_{nat}$). This means that if the neobiota added to a community are all very different from each other, the BNI (and Rao's Q) will keep increasing with newly added neobiota species. On the other hand, if neobiota all resemble each other, adding another species of neobiota to a community consisting already mainly of neobiota will create a decrease in BNI (and in Rao's Q)

**Overall, large mean trait differences between neobiota and natives tend to increase the hump shape, while high neobiota standard deviation tends to promote a monotonous increase in BNI.**

In scenario 4 (**Fig. S2.5 & S2.6**), we see that if neobiota are different from natives both in mean and in standard deviations, then both effects interact and the expected shape of the curve becomes more difficult to predict.

These BNI patterns follow closely the shape of Rao's Q. As a rule of thumb, the BNI follows a hump shaped curve when Rao's Q shows either a decreasing or a hump-shape curve. `

**On the other hand, the standardized BNI (BNIs) shows a strictly monotonous increase with increasing proportion of neobiota.**

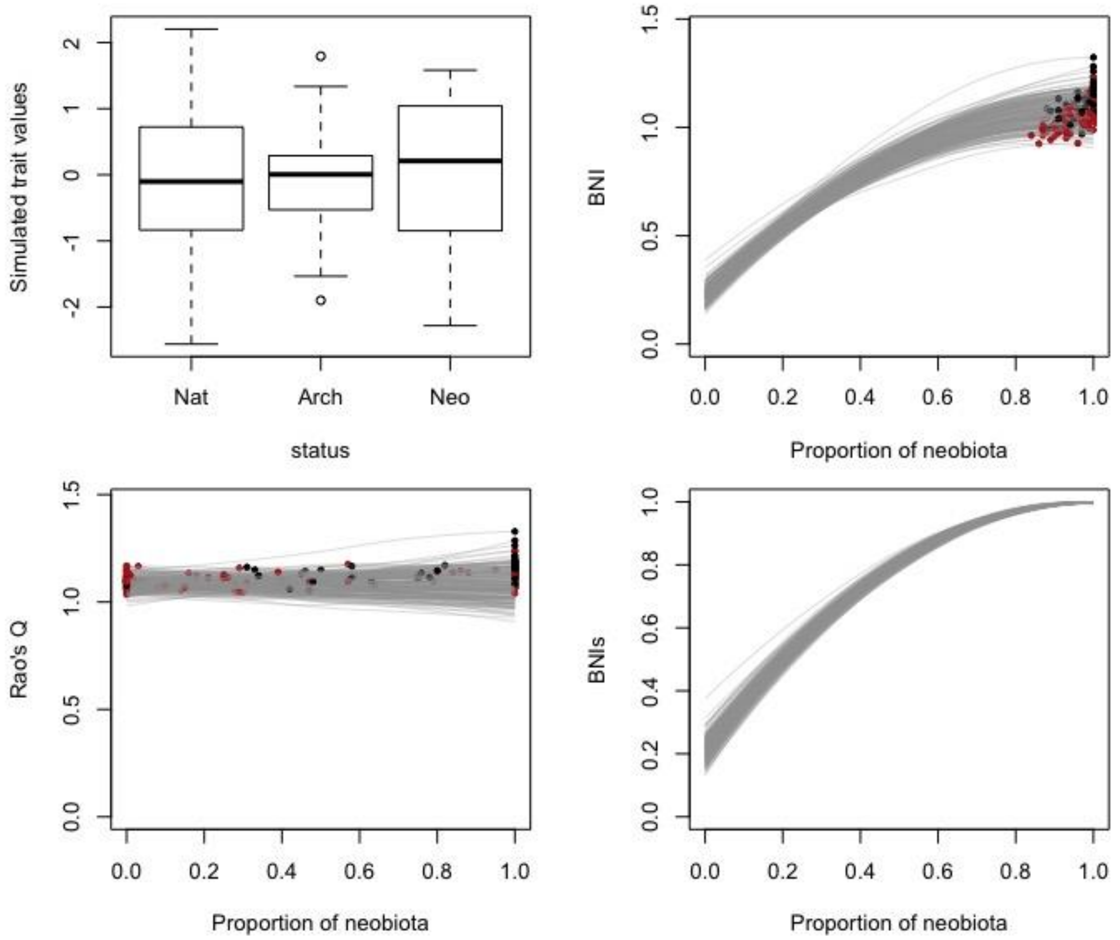


**Figure S2.1. Scenario 1: No difference in trait distributions between residents and neobiota.** All species trait values were randomly sampled from the same normal distribution of mean = 0 and SD = 1. Each grey line is a LOESS regression fitted to one of the 200 simulations. Dots represent the maximal value of each curve, with red dots representing simulations where SDneo < SDnat, and black dots SDneo ≥ SDnat. In this scenario, the SD were all very close to 1, but we can see that simulations (in red) where SDneo was randomly a little lower than SDnat, this small difference was enough to create a lower BNI maximum value. The upper left panel represents the trait T3 distribution in one example simulation of scenario 1 (Nat = natives, Arch = archaeobiota, Neo = neobiota).

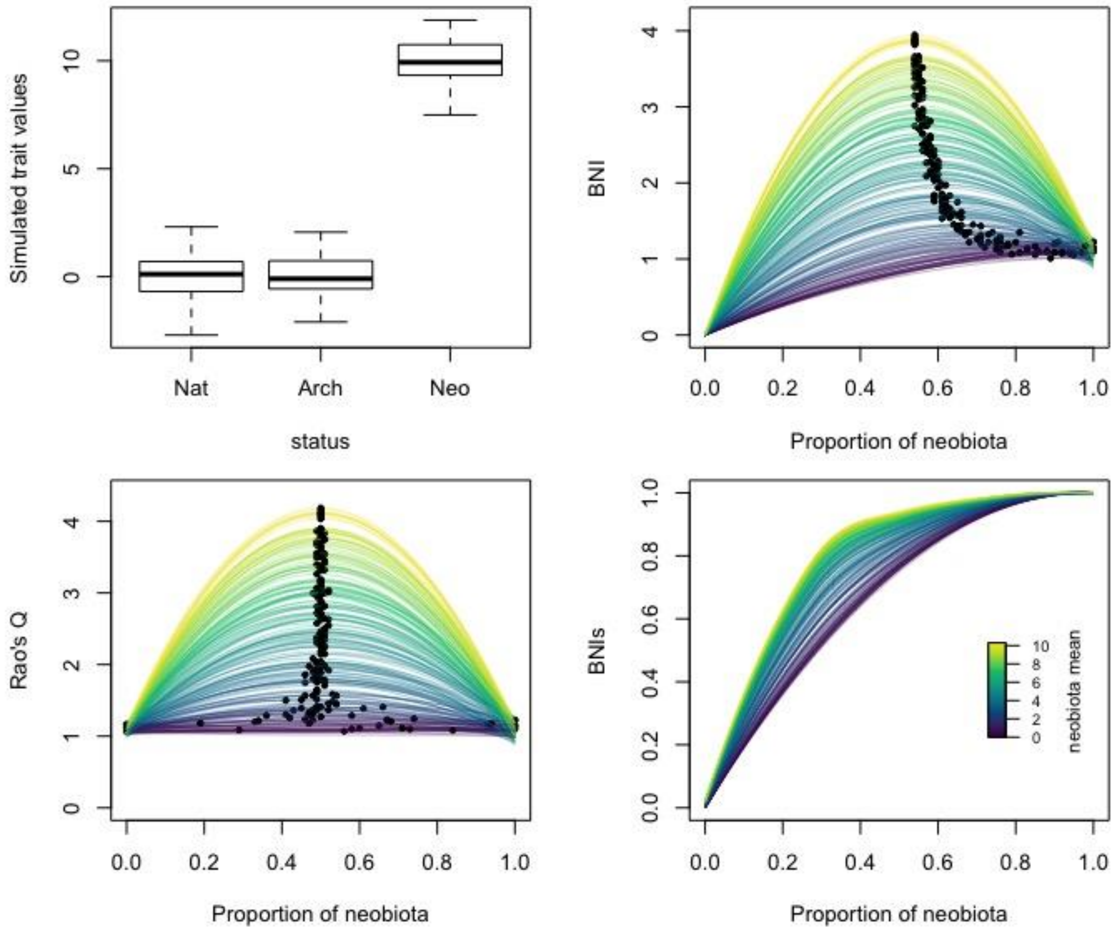


**Figure S2.2. Scenario 2: Neobiota differ on average from residents in at least one trait value**. Neophyte trait values are sampled from a normal distribution of mean = 0 to 10, and SD = 1. Natives and archaeobiota were all sampled from a normal distribution of mean = 0 and SD = 1. Colors represent the mean of the distribution from which neobiota are sampled, from 0 to 10 (in units of SD). Each line represents one of the 200 simulations. Dots represent the maximal value of each curve. The upper left panel represents an example of the trait T3 distribution in the most extreme simulation of scenario 2 (i.e. neobiota mean = 10; Nat = natives, Arch = archaeobiota, Neo = neobiota).

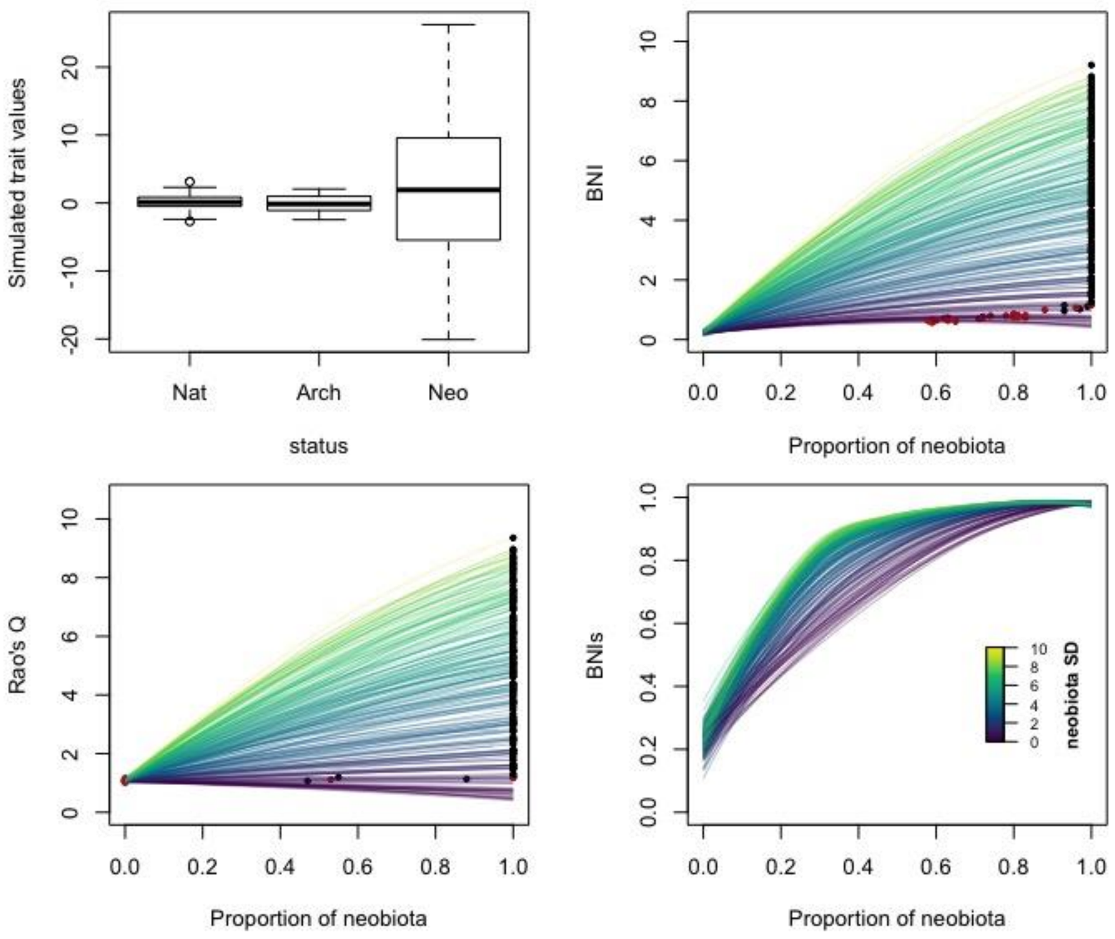


**Figure S2.3. Scenario 3a: Neophyte trait values occupy a wider range of trait values (i.e. higher standard deviation) than the residents.** Colors represent the standard deviation (SD) of the distribution from which neophyte trait values were sampled (mean = 0, SD from 0 to 10. Natives and archaeobiota were all sampled from a normal distribution of mean = 0 and SD = 1. Each line represents one of the 200 simulations. Dots represent the maximal value of each curve: red dots representing simulations where SDneo < SDnat; black dots SDneo ≥ SDnat. The upper left panel represents an example of the trait T3 distribution in the most extreme simulation of scenario 3 (i.e. neobiota SD = 10; Nat = natives, Arch = archaeobiota, Neo = neobiota).

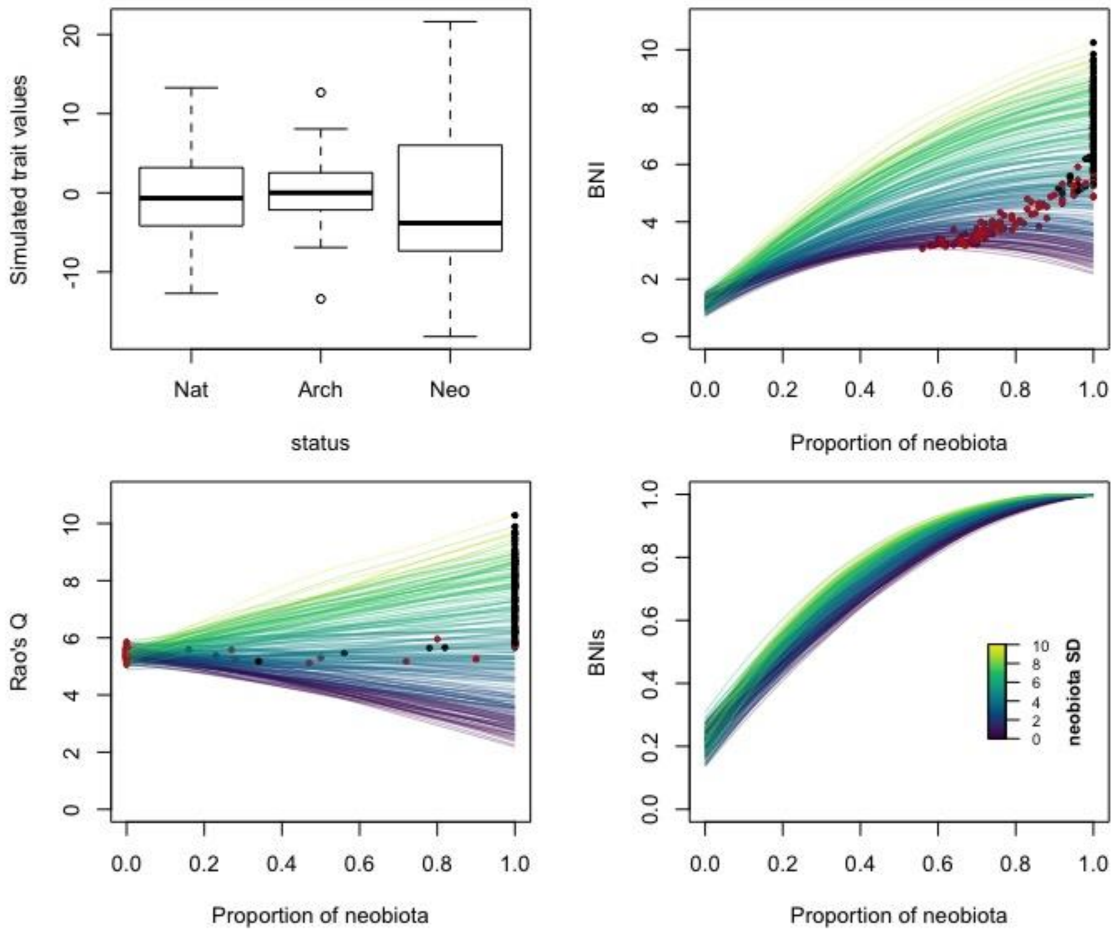


**Figure S2.4. Scenario 3b: Neobiota trait values occupy a different range of trait values (i.e. standard deviation) than the residents: from a lower to a higher range.** Colors represent the standard deviation (SD) of the distribution from which neophyte trait values were sampled (mean = 0, SD from 0 to 10. Natives and archaeobiota were all sampled from a normal distribution of mean = 0 and SD = 5. Each line represents one of the 200 simulations. Dots represent the maximal value of each curve: red dots representing simulations where SDneo < SDnat; black dots SDneo ≥ SDnat. The upper left panel represents an example of the trait T3 distribution in the most extreme simulation of scenario 3 (i.e. neobiota SD = 10; Nat = natives, Arch = archaeobiota, Neo = neobiota).

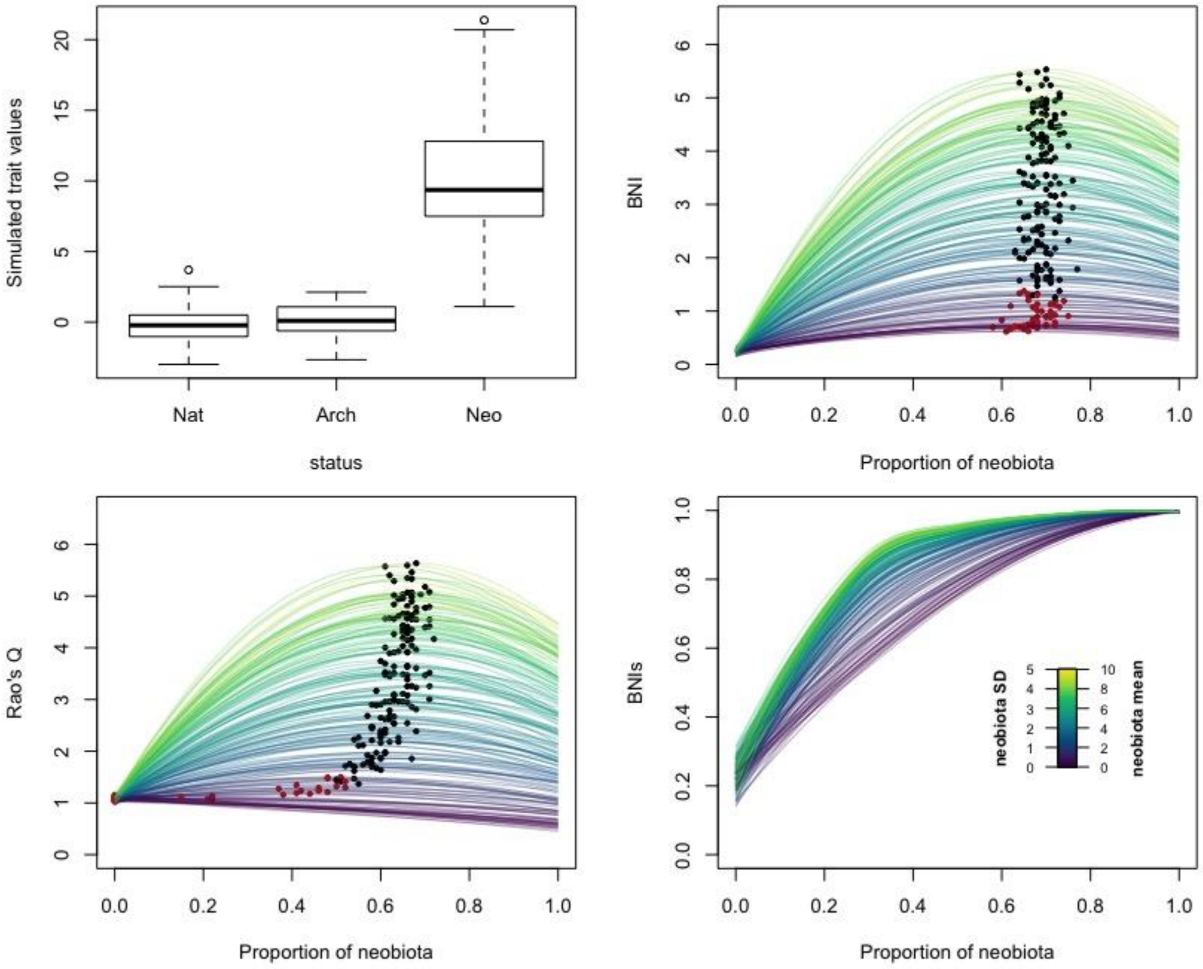


**Figure S2.5. Scenario 4a: Neobiota differ both on average and in variance from residents, with variance increasing with mean differences.** Neobiota were sampled from normal distributions of increasing mean (from 0 to 10) and increasing variance (from 0 to 5), as illustrated by the color gradient. Natives and archaeobiota remained sampled from a normal distribution of mean = 0 and SD = 1. Each line represents one of the 200 simulations. Dots represent the maximal value of each curve: red dots representing simulations where SDneo < SDnat; black dots SDneo ≥ SDnat. The upper left panel represents an example of the trait T3 distribution in the most extreme simulation of scenario 4 (i.e. neobiota mean = 10 and SD = 5; Nat = natives, Arch = archaeobiota, Neo = neobiota).

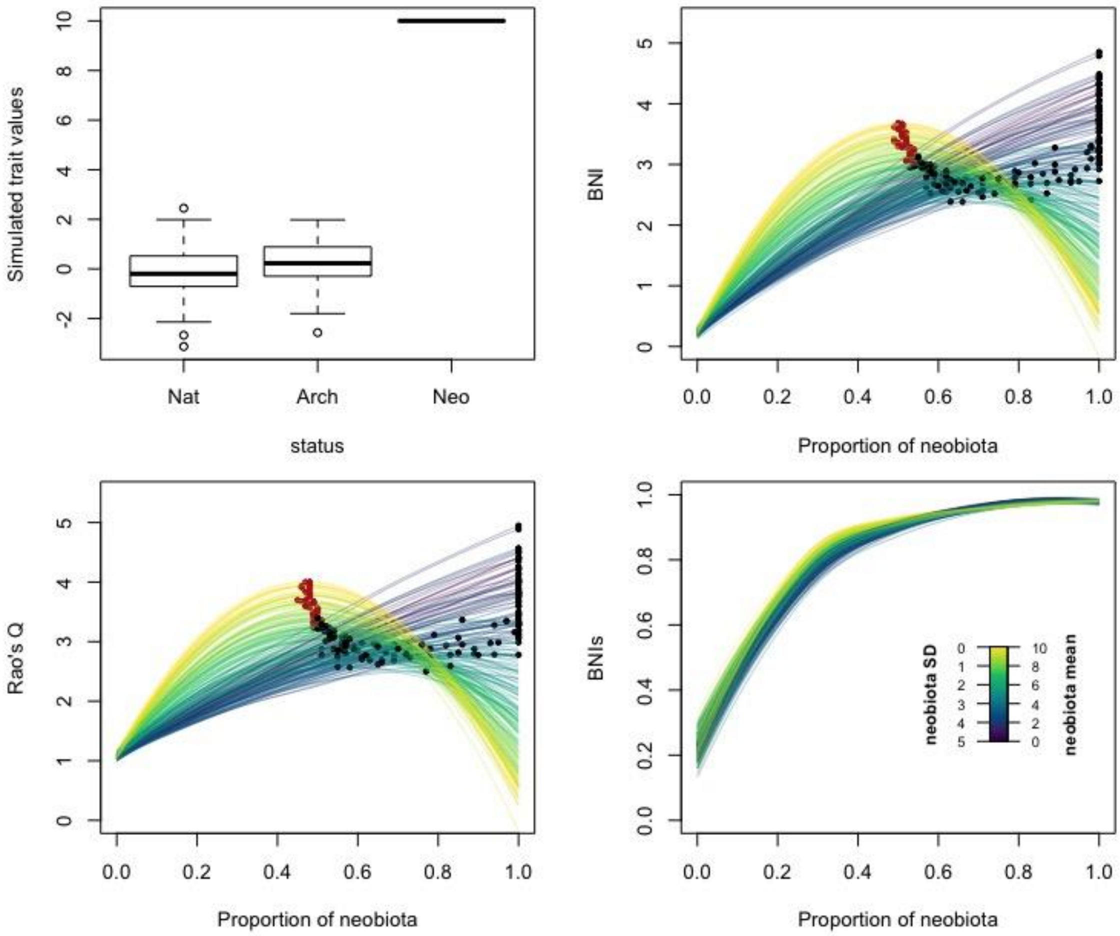


**Figure S2.6. Scenario 4b: Neobiota differ both on average and in variance from residents, with variance decreasing while mean differences increase.** Neobiota were sampled from normal distributions of increasing mean (from 0 to 10) and decreasing variance (from 5 to 0), as illustrated by the color gradient. Natives and archaeobiota remained sampled from a normal distribution of mean = 0 and SD = 1. Each line represents one of the 200 simulations. Dots represent the maximal value of each curve: red dots representing simulations where SDneo < SDnat; black dots SDneo ≥ SDnat. The upper left panel represents an example of the trait T3 distribution in the most extreme simulation of scenario 4 (i.e. neobiota mean = 10 and SD = 0; Nat = natives, Arch = archaeobiota, Neo = neobiota).
