## Supplementary material 3 - R code for "A multidimensional framework for measuring biotic novelty: How novel is a community?": FUNCTION BNI.calc.R

```
# BIOTIC NOVELTY INDEX 
# written by Conrad Schittko and Maud Bernard-Verdier, July 2019

#______________________________________________________________________________________________________________________#

# FUNCTION calculating an index of biotic novelty at the community level, based on species traits and years of residence.

## Inputs ## 
# com:          numeric community matrix of species abundance (sites x species)
# trait:        numeric matrix of trait values which will be used to calculate multitrait distances (species x traits)
# YSI:          numeric vector with "years since introduction" for each species in the region of interest (in years)
# dist.method:  character string of the name of a distance measure accepted by the function "dist" in package vegan

## Output : a list of elements comprising:
# $index 
      # $BNI:    numeric vector of BNI values per community site (cf. publication by Schittko et al., in prep)
      # $RaoQ:   numeric vector of Rao's quadratic entropy for each community
      # $BFI:    numeric vector of Biotic familiarity indices (BFI = RaoQ - BNI)
      # $prop.neo: proportion of neobiota in the community
# $trait.mat       trait matrix used to calculate distances
# $YSI             year since introduction of species                
# $rpi     vector of normalized residence time for each species
# $cij     matrix of temporal coexistence coefficient for each pair of species
# $t.dist  matrix of trait distances between pairs of species

#______________________________________________________________________________________________________________________#


BNI.calc <-
  function(com = abund,
           trait.mat = trait,
           dist.mat = NULL,
           YSI = status.data$years_since_introduction,
           dist.method = "gower") {
    require(FD)
    require(proxy)
    
    # Transform community matrix in relative abundances
    com <- com / rowSums(com)
    
    if (is.null(dist.mat)) {
    # Calculate pairwise trait distances "d(ij)"
    if (dist.method == "gower") {
      t.dist <- gowdis(x=trait.mat)
      t.dist <- as.matrix(t.dist)
    } else {
      t.dist <- dist(trait.mat, method = dist.method)
      t.dist <- as.matrix(t.dist)
    }
  } else {
    t.dist <- as.matrix(dist.mat)
  }
    
    # calculate the normalized time of residence of each species compared to the community "r'i"
    rpi <-  (YSI - min(YSI)) / (max(YSI) - min(YSI))
    
    # calculate pairwise temporal coefficients of coexistence "C(ij)"
    cij <- as.matrix(1 - dist(rpi, method = min))
    
    # combine matrices of trait distance with temporal coefficient into one weight matrix : "d(ij) x c(ij)"
    dxc <- t.dist * cij
    dxc <- as.matrix(dxc)
    
    # Calculate the BNI as a cross product of two matrices: community matrix x weight matrix
    BNI <- apply(com, 1, function(x)
      crossprod(x, dxc %*% x)) / 2
    
    # Calculate also Rao's quadratic entropy
    RaoQ <- apply(com, 1, function(x)
      crossprod(x, t.dist %*% x)) / 2
    
    # Calculate the BFI (Biotic familiarity component) as the complementary partition of BNI:
    BFI <- RaoQ - BNI
    
    # calculate proportion of neophytes
    neos <-  as.numeric(YSI<=520)
    prop.neo <- rowSums(ceiling(com)[, neos == 1]) / rowSums(ceiling(com))

    return(list(
      index = data.frame(BNI = BNI,
      RaoQ = RaoQ,
      BFI = BFI,
      prop.neo = prop.neo),
      trait.mat = trait.mat,
      YSI = YSI,
      rpi = rpi,
      cij = cij,
      t.dist = t.dist
    ))
  }
```
