## Supplementary material 3 - R code for "A multidimensional framework for measuring biotic novelty: How novel is a community?": FUNCTIONS BNI simulations.R

```
# Simulations to test and visualize the BNI
# written by Conrad Schittko and Maud Bernard-Verdier, Obtober 2019

#_________________________________________________________#
# This code comprises five functions, making use of the function BNI.calc:

# 1. simul.comm
# Generates a random sets of communities, species and traits, with predetermined proportion of alien species and trait differences between species

# 2. simul.BNI
# Wrapper function for simul.comm, to generate multiple set of random communities and calculate associated BNI, Rao and BNIs indices

# 3. plot.simBNI
# plotting function for a graphical representation of the output of simul.BNI

### Some utility functions:

# 4. add.stats
# basic utility function to add lm or glm statistics to plots

# 4. my.image.plot
# function "image.plot" in package "fields"
# slightly modified to add more plotting options 
#______________________________________________________#


#### 1. FUNCTION simul.comm : Community assembly simulations  ####

## Inputs ##
# nsp             total number of species in the speices pool
#
# nsites          number of community sites to generate
#
# mean.rich       mean species richness in communities
#
# simul.com      method for simulating community assembly, to choose from :
#   "random"              random species occurrences, random proportion of neobiota
#   "prop.neo"            communities have predetermined proportion of neobiota
#   "trait.filter"        communities are distributed in habitats selecting different trait means
#
# simul.trait     method for simulating trait values, to choose from :
#   "random"      trait values sampled from normal distribution with mean=O and SD=sim.SD for all species
#   "mean difference"    traits from normal distributions with different mean for neobiota
#   "variance difference" traits from normal distributions with different SD for neobiota
#   "both different"      traits from normal distributions with different mean and SD for neobiota
#
# p.status        numeric vector of proportions of natives, archaeobiota and neobiota in species pool.
#                 Default is: c(nat = 0.70, arch = 0.15, neo = 0.15). Must be of length 3.
#
# t.arrival       numeric vector providing the residence times (in years) of species
#                 for different introduction status: value for native, value for archeobiota,
#                 minimum for neobiota and maximum for neobiota. 
#                 Default is:  c(nat = 8518, arch = 8518, neo.min = 1, neo.max = 526)
#
# p.neo           numeric vector of proportions of neobiota in communities,
#                 which will be distributed randomly across "nb.sites" communites.
#                 Default is: c(0, 0.25, 0.5, 0.75, 1)
#
# deltaTrait      mean trait difference of neobiota for simul.trait = c("mean difference","both different")
#
# s.SD = 1        default standard deviation of trait distributions (including for natives)
#
# neoSD           Numeric value,  SD for neobiota traits when
#                 simul.trait = c("variance difference","both different").


## Output : a list of elements with results of one set of simulated communities
# species.data    simulated introduction status and trait data for the nsp species
# comMat          simulated community matrix of of dimension (nsites, nsp)
# hab             sequence of different habitat modelled (not used)
# mean.rich       mean simulated richness
# nspecies        number of simulated species in the pool
# nsites          number of sites (communities) simulated
# Simul.trait     trait scenario
# Simul.com       communitty assembly scenario
# propNeo         vector of proportion of neophytes for each habitat
# #
#_______________________________________________________________________

simul.comm <- function(nsp = 250,
                       nsites = 100,
                       mean.rich = 25 ,
                       simul.trait = c("random"),
                       simul.com = c("random"),
                       p.status = c(nat = 0.70, arch = 0.15, neo = 0.15),
                       t.arrival = c(nat = 8518, arch = 8518,
                                     neo.min = 1, neo.max = 526),
                       p.neo = c(0, 0.25, 0.5, 0.75, 1),
                       deltaTrait = 0.3,
                       s.SD = 1,
                       neo.SD = s.SD) {
  ## simulate species data
  
  species.data <- data.frame(species = paste("sp", 1:nsp, sep = ""))
  
  # assign species status (1 = native, 2 = archeobiota, 3 = neobiota), with 70% of natives
  species.data$status  <-
    sample(c(1, 2, 3), nsp, replace = T, prob = p.status)
  
  ## Assign years since introduction based on the status
  
  # Natives estimated to be resident since - 6500 AD
  species.data$YSI <- t.arrival[1]
  
  # archaeobiota introduced approx. -750 AD
  species.data$YSI[species.data$status == 2] <-  t.arrival[2]
  
  # Uniform distribution to assign introduction years of neobiota since 1492
  species.data$YSI[species.data$status == 3] <-
    ceiling(runif(
      sum(species.data$status == 3),
      min = t.arrival[3],
      max = t.arrival[4]
    ))
  
  
  # TRAIT SCENARIO A : no difference in traits between natives, archaeobiota and neobiota
  
  if (simul.trait == "random") {
    species.data$T1 = rnorm(nsp, mean = 0 , sd = s.SD)
    species.data$T2 = rnorm(nsp, mean = 0 , sd = s.SD)
    species.data$T3 =  rnorm(nsp, mean = 0 , sd = s.SD)
  }
  
  # TRAIT SCENARIO B : mean difference in trait between natives and neobiota, same variance
  
  if (simul.trait == "mean difference") {
    neo <- species.data$status == 3  ## Define neobiota
    species.data$T1[!neo] = rnorm(sum(!neo), mean = 0 , sd = s.SD)
    species.data$T2[!neo] = rnorm(sum(!neo), mean = 0 , sd = s.SD)
    species.data$T3[!neo] =  rnorm(sum(!neo), mean = 0 , sd = s.SD)
    
    # generate different trait values for neobiota, only for two out of three traits:
    species.data$T1[neo] = rnorm(sum(neo), mean = deltaTrait , sd = s.SD)
    species.data$T2[neo] = rnorm(sum(neo), mean = deltaTrait , sd = s.SD)
    species.data$T3[neo] =  rnorm(sum(neo), mean = 0 , sd = s.SD)
  }
  
  # TRAIT SCENARIO C :difference in trait variance between natives and neobiota, same mean
  
  if (simul.trait == "variance difference") {
    neo <- species.data$status == 3  ## Define neobiota
    species.data$T1[!neo] = rnorm(sum(!neo), mean = 0 , sd = s.SD)
    species.data$T2[!neo] = rnorm(sum(!neo), mean = 0 , sd = s.SD)
    species.data$T3[!neo] =  rnorm(sum(!neo), mean = 0 , sd = s.SD)
    
    
    # generate different trait values for neobiota, only for two out of three traits:
    species.data$T1[neo] = rnorm(sum(neo), mean = 0, sd = neo.SD)
    species.data$T2[neo] = rnorm(sum(neo), mean = 0, sd = s.SD)
    species.data$T3[neo] =  rnorm(sum(neo), mean = 0, sd = s.SD)
  }
  
  
  # TRAIT SCENARIO D : difference in trait mean and variance between natives and neobiota
  
  if (simul.trait == "both different") {
    neo <- species.data$status == 3  ## Define neobiota
    species.data$T1[!neo] = rnorm(sum(!neo), mean = 0 , sd = s.SD)
    species.data$T2[!neo] = rnorm(sum(!neo), mean = 0 , sd = s.SD)
    species.data$T3[!neo] =  rnorm(sum(!neo), mean = 0 , sd = s.SD)
    
    
    # generate different trait values for neobiota, only for one out of three traits:
    species.data$T1[neo] = rnorm(sum(neo), mean = deltaTrait, sd = neo.SD)
    species.data$T2[neo] = rnorm(sum(neo), mean = deltaTrait, sd = s.SD)
    species.data$T3[neo] =  rnorm(sum(neo), mean = 0, sd = s.SD)
  }
  
  #### COMMUNITY ASSEMBLY
  
  # ASSEMBLY 0: total random
  if (simul.com == "random") {
    # define species richness (SR) for nsites site-level communities, from nsp species
    SR <-  c(rpois(nsites, lambda = mean.rich))
    # ransomly permute site SR
    SR <- sample(SR)
    
    # assign a habitat randomly to communities, from 4 categories
    hab <- sample(rep(c(1, 2, 3, 4), 14))
    
    # assemble communities randomly:
    comMat <- t(sapply(SR, function(x) {
      r <- rep(0, nsp)
      r [sample(1:nsp, x)] <- 1
      return(r)
    }))
    rownames(comMat) <- paste("site", 1:nsites, sep = "")
    colnames(comMat) <- species.data$species
  }
  
  # ASSEMBLY 2: some habitats have more and more neobiota
  if (simul.com == "prop.neo") {
    
    # define species richness (SR) for nsites site-level communities, from nsp species
    SR <-  c(rpois(nsites, lambda = mean.rich))
    
    # Proportion of neobiota across the 5 habitats
    prop.neo <- p.neo
    
    # assign habitats uniformly
    hab <-
      sample(rep_len(x = 1:length(p.neo) , length.out = nsites))
    
    # assemble communities randomly based on expected neobiota proportions:
    comMat <- t(sapply(1:nsites, function(i) {
      r <- rep(0, nsp)
      
      Sneo <- ceiling(SR[i] * prop.neo[hab[i]])
      Snat <- ceiling(SR[i] * (1 - prop.neo[hab[i]]))
      
      if (Sneo > sum(species.data$status == 3))
        Sneo = sum(species.data$status == 3)
      if (Snat > sum(!species.data$status == 3))
        Snat = sum(!species.data$status == 3)
      
      # select the natives and archaeobiota
      r [sample(which(!species.data$status == 3), Snat)] <- 1
      
      # select the aliens
      r [sample(which(species.data$status == 3), Sneo)] <- 1
      
      return(r)
    }))
    rownames(comMat) <- paste("site", 1:nsites, sep = "")
    colnames(comMat) <- species.data$species
  }
  
  
  
  # ASSEMBLY 3: predetermined mixes of native, archaeobiota and neobiota
  if (simul.com == "fixed.mix") {
   
    # define species richness (SR) for nsites site-level communities, from nsp species
    SR <-  c(rpois(nsites, lambda = mean.rich))
    
    # Proportion of neobiota and archaeobiota in the different habitat types
    mixes <- data.frame(h1 = c(1,0,0),
                   h2 = c(0.5,0.5,0),
                   h3 = c(0,1,0),
                   h4 = c(0.5,0, 0.5),
                   h5 = c(1/3,1/3,1/3),
                   h6 = c(0,0.5,0.5),
                   h7 = c(0,0,1))
    
    # assign habitats uniformly
    hab <-
      sample(rep_len(x = 1:length(mixes) , length.out = nsites))
    
    # assemble communities randomly based on expected neobiota proportions:
    comMat <- t(sapply(1:nsites, function(i) {
      r <- rep(0, nsp)
      
      Sneo <- ceiling(SR[i] * mixes[3,hab[i]])
      Sarc <- ceiling(SR[i] * mixes[2,hab[i]])
      Snat <-  ceiling(SR[i] * mixes[1,hab[i]])
      
      if (Sneo > sum(species.data$status == 3))
        Sneo = sum(species.data$status == 3)
      if (Snat > sum(species.data$status == 1))
        Snat = sum(species.data$status == 1)
      if (Sarc > sum(species.data$status == 2))
        Sarc = sum(species.data$status == 2)
      
      # select the natives 
      r [sample(which(species.data$status == 1), Snat)] <- 1
      
      # select the archaeobiota
      r [sample(which(species.data$status == 2), Sarc)] <- 1
      
      # select the aliens
      r [sample(which(species.data$status == 3), Sneo)] <- 1
      
      return(r)
    }))
    rownames(comMat) <- paste("site", 1:nsites, sep = "")
    colnames(comMat) <- species.data$species
  }
  
  propNeo <-
    rowSums(comMat[, species.data$status == 3]) / rowSums(comMat)
  
  simul <-
    list(
      species.data = species.data,
      comMat = comMat ,
      hab = hab ,
      mean.rich = mean.rich,
      nspecies = nsp,
      nsites = nsites,
      Simul.trait = simul.trait ,
      Simul.com = simul.com ,
      propNeo = propNeo
    )
  
  return(simul)
}

#_______________________________________________________________________
#### 2. FUNCTION simul.BNI:  Apply simulations and calculate BNI ####
# This funciton is a wrapper which applies functions simul.comm and BNI.calc for a range of simulation conditions, such as increasing trait differences between natives and neobiota.
# The number of simulations can be increased with "nreps" if results of simulations are too variable (this will take more time).
#
## INPUTS ##
#
## Inputs for individual community simulations:
# simulation.trait   character string. input simul.trait for function simul.comm
# simulation.comm    character string. input simul.com for function simul.comm
# nb.species         number of species to simulate in species pool
# nb.sites           number of communities/sites to simulate
# mean.richness      mean richness of the simulated communities (all have the same SR expectation)
# proportion.status  numeric vector of proportions of natives, archaeobiota and neobiota in species pool.
# proportion.neo     numeric vector of proportions of neobiota in communities,
#                    which will be distributed randomly across "nb.sites" communites.
# time.arrival       numeric vector defining input t.arrival for function simul.comm
# sim.SD             baseline standard deviation of trait distributions
# 
## Inputs for the BNI calculation:
# distance.method    character string for the input "dist.meth" in funciton BNI.calc.
#                    Must correspond either to one of the methods in function "dist" 
#                    of package "stats", or be "gower" for implementing 
#                    the function "gowdist" from package "FD".
#
## Inputs for controlling the sequence of repeated simulations:
# nreps= 2           number of repetitions of the sequence of nb.sites simulations
# maxDT              maximum mean trait difference to generate across the sequence of simulations
# maxDSD             maximum difference in variance to generate across the sequence of simulations
# sd.trend           method for ordering sequences of trait mean and variance for neophtyes.
#                    Default is positive correlation, with SD increasing with mean.
#                    May be set to "random", or "inverse".
#
## OUTPUT
# The function returns a list of simulation results, of length the number of repetitions 'nreps'
# Each repetition has the following structure:
# $index:   data.frame of n.site rows, with :
#             $BNI
#             $RaoQ
#             $BFI (Biotic index of Familiarity: BFI =  Rao's Q - BNI) 
#             $prop.neo: proportion or neophytes simulated
#             $hab : habitat type (1 to 5: defining % of neophyte expected)
#
# $species.data   data.frame of values for the simulated n.species : 
#                   $status:  (1:native, 2:archaeo, 3: neobiota),
#                   $YSI : years since introduction,
#                   $T1 to $T3 :  the three simulated traits 
#
# $simul.trait      scenario chosen for the trait simulation 
# $sd.trend         option chosen for the trait simulation when both mean and #                   SD of neobiota are different from natives 
#_______________________________________________________________________
simul.BNI <-
  function(simulation.trait = "mean difference",
           simulation.comm = "prop.neo",
           distance.method =  "Euclidean",
           nreps = 2 ,
           nb.species = 250,
           nb.sites = 100,
           mean.richness = 25,
           proportion.status = c(nat = 0.70, arch = 0.15, neo = 0.15),
           proportion.neo = c(0, 0.25, 0.5, 0.75, 1),
           maxDT = 10,
           maxDSD = 10,
           sim.SD = 1,
           sd.trend = "correlated",
           time.arrival = c(nat = 8518, arch = 2768,
                            neo.min = 1, neo.max = 100))
  {
    # sequence of 20 levels of trait differences for neobiota, multiplied by nreps.
    seqDT <- rep(seq(0, maxDT, maxDT / 19), nreps)
    seqDSD <- rep(seq(0, maxDSD, maxDSD / (19)), nreps)
    seq.simSD <- rep(sim.SD, nreps*20)
    
    if (sd.trend == "all increase") {
    seq.simSD <- rep(seq(0, sim.SD, sim.SD / (19)), nreps)
    }
    
    if (sd.trend == "random")
      seqDSD = sample(seqDSD)
    
    if (sd.trend == "inverse")
      seqDSD = seqDSD[(20 * nreps):1]
    
    sim_list <- sapply(
      1:(20 * nreps),
      FUN = function(i)
      {
        sim <- simul.comm(
          simul.trait = simulation.trait ,
          simul.com = simulation.comm,
          nsp = nb.species,
          nsites = nb.sites,
          mean.rich = mean.richness ,
          p.status = proportion.status,
          p.neo = proportion.neo,
          deltaTrait = seqDT[i],
          s.SD = seq.simSD[i],
          neo.SD = seqDSD[i],
          t.arrival = time.arrival
        )
        
        bni <- BNI.calc(
          com = sim$comMat,
          trait.mat = sim$species.data[, c("T1", "T2", "T3")],
          YSI = sim$species.data$YSI ,
          dist.method = distance.method
        )
        
        bni$index$hab <- sim$hab
        
        return(
          list(
            index = bni$index,
            species.data = sim$species.data,
            simul.trait = simulation.trait,
            sd.trend = sd.trend
          )
        )
      },
      simplify = FALSE
    )
    return(sim_list)
  }

#_______________________________________________________________________
#### 3. FUNCTION plot.simBNI :  Graphical output of BNI simulation ####

## Inputs ##
# sim_list  an output object from simul.BNI
# type      vector of strings to select which combination 
#           of graphs to plot. Can be a combination of 
#           the following: c("Full", "BNI", "BNIs", "Rao").
#           Default to "Full".
# color.option     a character string indicating if the color gradient
#                  should vary with the different in trait mean or 
#                  SD of the neobiota in each simulation. 
#                  option = NULL colors will follow the mean variations.
# color.bar        logical: whether to plot a color bar legend
#
# which.bar        a character string indicating if two color 
#                  bars (SD and mean) should be represented together
#                  or in reverse order. 
#                  Default is NULL, only one bar will be plotted. 
#                 "inverse" is in reverse order. 
#                 "together" is in the same order.
#
# max.point        logical, indicating whether maximum points 
#                  should be drawn on the curves. Default is TRUE.
#
# diff.col.max    logical, whether different colors should 
#                 be used to color the maximum points. Dafault is FALSE.
#                 If TRUE: red = SDneo < SDnat, black = SDneo >= SDnat
#                 
#
## Output : a four panel plot
#_______________________________________________________________________
plot.BNIsim <-
  function(sim_list,
           type = c("Full", "BNI", "BNIs", "Rao", "Traits"),
           xAxis = c("prop.neo", "mixes"),
           color.option = c("mean"),
           color.bar = TRUE, 
           which.bar = NULL,
           max.point = TRUE, 
           diff.col.max = TRUE) {
    
    require(fields)
  
# Define color scheme  
    mean.seq = unlist(lapply(
      sim_list,
      FUN = function(x)
        mean(x$species.data$T1[x$species.data$status == 3])
    ))
    sd.seq = unlist(lapply(
      sim_list,
      FUN = function(x)
        sd(x$species.data$T1[x$species.data$status == 3])
    ))
    
    sd.nat.seq = unlist(lapply(
      sim_list,
      FUN = function(x)
        sd(x$species.data$T1[x$species.data$status !=3])
    ))
    
    
    allsd.seq = unlist(lapply(
      sim_list,
      FUN = function(x)
        sd(x$species.data$T1)
    ))
    
    
    col.seq = rep("#A1A1A1", length(sim_list))
    
    if (color.option == "mean") {
      sq <- floor((mean.seq - min(mean.seq)) / sd(mean.seq) *100) + 1
      col.seq = hcl.colors(max(sq))[sq]
    }
    
    if (color.option == "SD") {
      sq <- floor((sd.seq - min(sd.seq)) / sd(sd.seq) * 100) + 1
      col.seq = hcl.colors(max(sq))[sq]
    } 
    
    if (color.option == "allSD") {
      sq <- floor((allsd.seq - min(allsd.seq)) / sd(allsd.seq) * 100) + 1
      col.seq = hcl.colors(max(sq))[sq]
    } 
    
    col.seq = paste(col.seq, "50", sep = "")
    
    if (xAxis[1] != "mixes") {
    ## Plot only the trait values simulated
    if (type[1] == "Traits") {
      par(
        mfrow = c(1, 5),
        mar = c(1,0, 1, 0),
        oma = c(1, 4, 1, 1)
      )
      for (i in c(1, 5, 10, 15, 20)) {
        sim <- sim_list[[i]]
        boxplot(
          T1 ~ status,
          sim$species.data,
          names = c("Nat", "Arch", "Neo"),
          ylim = c(-15, 15), 
          yaxt = "n"
        )
        if (i == 1) axis(2)
      }
    }
    
    ## BNI plots
    if (type[1] != "Traits") {
    
      # Select the plots to include
      nb.fig = length(type)
      if (type[1] == "Full")
        nb.fig = 4
      
      if (nb.fig == 1)
        par(
          mfrow = c(1, 1),
          mar = c(4, 4, 1, 2),
          oma = c(1, 1, 1, 1)
        )
      if (nb.fig == 2)
        par(
          mfrow = c(1, 2),
          mar = c(4, 4, 1, 2),
          oma = c(1, 1, 1, 1)
        )
      if (nb.fig == 3)
        par(
          mfrow = c(1, 3),
          mar = c(4, 4, 1, 2),
          oma = c(1, 1, 1, 1)
        )
      if (nb.fig == 4)
        par(
          mfrow = c(2, 2),
          mar = c(4, 4, 1, 2),
          oma = c(1, 1, 1, 1)
        )
      
      
      ## Illustrate max simulated trait differences
      if ("Full" %in%  type) {
        sim_max <- sim_list[[length(sim_list)]]
        boxplot(
          T1 ~ status,
          sim_max$species.data,
          names = c("Nat", "Arch", "Neo"),
          ylab = "Simulated trait values"
        )
      }
      
      ## BNI
      if ("BNI" %in% type | "Full" %in% type) {
        nd = data.frame(prop.neo = seq(0, 1, 0.01))
        y.max <-
          max(unlist(lapply(
            sim_list,
            FUN = function(x)
              max(x$index$BNI)
          )))
        
        max(c(sim_list[[1]]$index$BNI, sim_list[[20]]$index$BNI))
        plot(c(0, 1), c(0,  y.max),
             type = "n",  ann = F)
        title(xlab = "Proportion of neobiota", ylab = "BNI")
        
        
        for (i in 1:length(sim_list)) {
          lo <- loess(BNI ~ prop.neo, data = sim_list[[i]]$index, span = 0.85)
          plo <-
            predict(lo, newdata = data.frame(prop.neo = seq(0, 1, 0.01)), se = TRUE)
          lines(nd$prop.neo, plo$fit, col = col.seq[i])
          
          col.max = "black"
          if (diff.col.max) {
          col.max = "firebrick"
          if (sd.seq[i] > sd.nat.seq[i]) col.max <- "black"
          }
          
          
          if(max.point){
          mi <- which(plo$fit== max(plo$fit))
          points(seq(0, 1, 0.01)[mi],
                 plo$fit[mi],
                 col = col.max, cex = 0.8, pch = 20 )
          }
        }
      }
      
      ## Rao
      if ("Rao" %in% type | "Full" %in% type) {
        nd = data.frame(prop.neo = seq(0, 1, 0.01))
        
        y.max <-
          max(unlist(lapply(
            sim_list,
            FUN = function(x)
              max(x$index$RaoQ)
          )))
        
        plot(c(0, 1), c(0, y.max),
             type = "n",  ann = F)
        title(xlab = "Proportion of neobiota", ylab = "Rao's Q")
        
        for (i in 1:length(sim_list)) {
          lo <- loess(RaoQ ~ prop.neo, data = sim_list[[i]]$index, span = 0.85)
          plo <- predict(lo, newdata = data.frame(prop.neo = seq(0, 1, 0.01)), se = TRUE)
          lines(nd$prop.neo, plo$fit, col = col.seq[i])
          
          col.max = "black"
          if (diff.col.max) {
            col.max = "firebrick"
            if (sd.seq[i] > sd.nat.seq[i]) col.max <- "black"
          }
          
             
          if(max.point){
            mi <- which(plo$fit== max(plo$fit))
            points(seq(0, 1, 0.01)[mi],
                   plo$fit[mi],
                   col = col.max, cex = 0.8, pch = 20 )
          }
          
        }
      }
      
      
      
      
      ## BNI/Rao
      if ("BNIs" %in% type | "Full" %in% type) {
        nd = data.frame(prop.neo = seq(0, 1, 0.01))
        plot(c(0, 1), c(0, 1),
             type = "n",  ann = F)
        title(xlab = "Proportion of neobiota", ylab = "BNIs")
        
        for (i in 1:length(sim_list)) {
          BNIs <- (sim_list[[i]]$index$BNI / sim_list[[i]]$index$RaoQ)
          lo <- loess(BNIs ~ prop.neo, data = sim_list[[i]]$index, span = 0.85)
          plo <-
            predict(lo, newdata = data.frame(prop.neo = seq(0, 1, 0.01)), se = TRUE)
          lines(nd$prop.neo, plo$fit, col = col.seq[i])
        }
      }
      
      ## Draw legend bar
      
      if (color.bar) {
        if (color.option == "mean" & is.null(which.bar)) {
          sd.seq = unlist(lapply(
            sim_list,
            FUN = function(x)
              sd(x$species.data$T1[x$species.data$status == 3])
          ))
          my.image.plot(
            legend.only = TRUE,
            zlim = c(0, max(mean.seq)),
            col = hcl.colors(200),
            smallplot = c(0.70, 0.73, 0.4, 0.6),
            legend.lab = "neobiota mean",
            legend.line = 2,
            legend.cex = 0.7,
            border = "white",
            axis.args = list(cex.axis = 0.7)
          )
        }
        
        if (color.option == "SD" &  is.null(which.bar)) {
          my.image.plot(
            legend.only = TRUE,
            zlim = c(0, floor(max(sd.seq))),
            col = hcl.colors(200),
            smallplot = c(0.70, 0.73, 0.4, 0.6),
            legend.lab = "neobiota SD",
            legend.line = 2,
            legend.cex = 0.7,
            border = "white",
            axis.args = list(cex.axis = 0.7)
          )
        }
        
        if (!is.null(which.bar)) {
          my.image.plot(
            legend.only = TRUE,
            zlim = c(0, max(mean.seq)),
            col = hcl.colors(200),
            smallplot = c(0.70, 0.73, 0.4, 0.6),
            legend.lab = "neobiota mean",
            legend.line = 2,
            legend.cex = 0.7,
            axis.at = seq(0, max(mean.seq), by = 2),
            axis.lab = seq(0, max(mean.seq), by = 2),
            border = "white",
            axis.args = list(cex.axis = 0.7)
          )
          
          my.image.plot(
            legend.only = TRUE,
            col = hcl.colors(200),
            zlim = c(0, floor(max(sd.seq))),
            smallplot = c(0.70, 0.73, 0.4, 0.6),
            border = "white",
            axis.args = list( cex.axis = 0.7),
            axis.lab = (if (which.bar == "inverse") {
              seq(floor(max(sd.seq)),0, by = -1)
              } else {
                seq(0, floor(max(sd.seq)), by = 1)
                }),
            axis.at = seq(0,floor(max(sd.seq)), by = 1),
            axis.side = 2,
            legend.args=list( text = "neobiota SD", side=2, line=2, cex=0.7)
          )
        }
        
      }
    }
    }
    
    ## Plot by mixes not prop neobiota
    if (xAxis[1] == "mixes") {
      
      # Select the plots to include
      nb.fig = length(type)
      if (type[1] == "Full")
        nb.fig = 4
      
      if (nb.fig == 1)
        par(
          mfrow = c(1, 1),
          mar = c(6, 4, 1, 2),
          oma = c(1, 1, 1, 1)
        )
      if (nb.fig == 2)
        par(
          mfrow = c(1, 2),
          mar = c(6, 4, 1, 2),
          oma = c(1, 1, 1, 1)
        )
      if (nb.fig == 3)
        par(
          mfrow = c(1, 3),
          mar = c(6, 4, 1, 2),
          oma = c(1, 1, 1, 1)
        )
      if (nb.fig == 4)
        par(
          mfrow = c(2, 2),
          mar = c(6, 4, 1, 2),
          oma = c(1, 1, 1, 1)
        )
      
      
      ## Illustrate max simulated trait differences
      if ("Full" %in%  type) {
        sim_max <- sim_list[[length(sim_list)]]
        boxplot(
          T1 ~ status,
          sim_max$species.data,
          names = c("Nat", "Arch", "Neo"),
          ylab = "Simulated trait values"
        )
      }
      
      ## BNI
      if ("BNI" %in% type | "Full" %in% type) {
        nd = data.frame(1:7)
        y.max <-
          max(unlist(lapply(
            sim_list,
            FUN = function(x)
              max(x$index$BNI)
          )))
        
        plot(c(1, 7), c(0,  y.max),
             type = "n",  ann = F, xaxt="n")
        title(xlab = "", ylab = "BNI")
        axis(1, at = 1:7, 
             labels = c("Natives only", "Arch + Nat","archaeobiota only",
                        "Nat + Neo", "Nat + Arch + Neo",
                        "Arch + Neo","neobiota only"),
             las = 2, cex.axis = 0.7)
        
        for (i in 1:length(sim_list)) {
          points(BNI ~ hab, data = sim_list[[i]]$index, col = col.seq[i])
        }
      }
      
      ## Rao
      if ("Rao" %in% type | "Full" %in% type) {
        nd = data.frame(prop.neo = seq(0, 1, 0.01))
        y.max <-
          max(unlist(lapply(
            sim_list,
            FUN = function(x)
              max(x$index$RaoQ)
          )))
        
        plot(c(1, 7), c(0,  y.max),
             type = "n",  ann = F, xaxt="n")
        title(xlab = "", ylab = "Rao's Q")
        axis(1, at = 1:7, 
             labels = c("Natives only", "Arch + Nat","archaeobiota only",
                        "Nat + Neo", "Nat + Arch + Neo",
                        "Arch + Neo","neobiota only"),
             las = 2, cex.axis = 0.7)
        
        for (i in 1:length(sim_list)) {
          points(RaoQ ~ hab, data = sim_list[[i]]$index,col = col.seq[i])
        }
      }
      
      ## BNI/Rao
      if ("BNIs" %in% type | "Full" %in% type) {
        
        plot(c(1, 7), c(0,  1),
             type = "n",  ann = F, xaxt="n")
        title(xlab = "", ylab = "BNIs")
        axis(1, at = 1:7, 
             labels = c("Natives only", "Arch + Nat","archaeobiota only",
                        "Nat + Neo", "Nat + Arch + Neo",
                        "Arch + Neo","neobiota only"),
             las = 2, cex.axis = 0.7)
        
        for (i in 1:length(sim_list)) {
          BNIs <- (sim_list[[i]]$index$BNI / sim_list[[i]]$index$RaoQ)
          points(BNIs ~ hab, data = sim_list[[i]]$index, col = col.seq[i])
        }     
      }
      
      ## Draw legend bar
      
      if (color.bar) {
        if (color.option == "mean") {
          sd.seq = unlist(lapply(
            sim_list,
            FUN = function(x)
              sd(x$species.data$T1[x$species.data$status == 3])
          ))
          my.image.plot(
            legend.only = TRUE,
            zlim = c(0, max(mean.seq)),
            col = hcl.colors(200),
            smallplot = c(0.70, 0.73, 0.4, 0.6),
            legend.lab = "neobiota mean",
            legend.line = 2,
            legend.cex = 0.7,
            border = "white",
            axis.args = list(cex.axis = 0.7)
          )
        }
        
        
        if (color.option == "SD") {
          my.image.plot(
            legend.only = TRUE,
            zlim = c(0,10),
            col = hcl.colors(200),
            smallplot = c(0.70, 0.73, 0.4, 0.6),
            legend.lab = "neobiota SD",
            legend.line = 2,
            legend.cex = 0.7,
            border = "white",
            axis.args = list(cex.axis = 0.7)
          )
        }
        
        if (color.option == "allSD") {
          my.image.plot(
            legend.only = TRUE,
            zlim = c(0, max(allsd.seq)),
            col = hcl.colors(200),
            smallplot = c(0.20, 0.23, 0.7, 0.9),
            legend.lab = "All species SD",
            legend.line = 2,
            legend.cex = 0.7,
            border = "white",
            axis.args = list(cex.axis = 0.7)
          )
        }
      }
    }
    
  }


## ADDITIONAL USEFUL FUNCTIONS FOR PLOTTING ####

# Function adding simple statistics of lm or glm to a basic plot:
add.stats <- function(f=NULL, formula = NULL, data= NULL) {
  if (is.null(f)) {
    f <- lm(as.formula(formula), data)
  }
  r2 = round(summary(f)$r.squared,2)
  p = anova(f)[1,5]
  mtext(3, text = paste(round(r2,2), p2star(p), sep = ""), adj = 1, cex = 0.7)
  if (p < 0.05) abline(f)
}


## function  'image.plot' from package 'fields',
## modified for a more flexible scale bar legend

my.image.plot <- function (..., add = FALSE, breaks = NULL, nlevel = 64, col = NULL, 
                           horizontal = FALSE, legend.shrink = 0.9, legend.width = 1.2, 
                           legend.mar = ifelse(horizontal, 3.1, 5.1), legend.lab = NULL, 
                           legend.line = 2, graphics.reset = FALSE, bigplot = NULL, 
                           smallplot = NULL, legend.only = FALSE, lab.breaks = NULL, 
                           axis.args = NULL, legend.args = NULL, legend.cex = 1, midpoint = FALSE, 
                           border = NA, lwd = 1, verbose = FALSE,
                           axis.side = NULL, axis.at = NULL, axis.lab = NULL) 
{
  old.par <- par(no.readonly = TRUE)
  if (is.null(col)) {
    col <- tim.colors(nlevel)
  }
  else {
    nlevel <- length(col)
  }
  info <- imagePlotInfo(..., breaks = breaks, nlevel = nlevel)
  breaks <- info$breaks
  if (verbose) {
    print(info)
  }
  if (add) {
    big.plot <- old.par$plt
  }
  if (legend.only) {
    graphics.reset <- TRUE
  }
  if (is.null(legend.mar)) {
    legend.mar <- ifelse(horizontal, 3.1, 5.1)
  }
  temp <- imageplot.setup(add = add, legend.shrink = legend.shrink, 
                          legend.width = legend.width, legend.mar = legend.mar, 
                          horizontal = horizontal, bigplot = bigplot, smallplot = smallplot)
  smallplot <- temp$smallplot
  bigplot <- temp$bigplot
  if (!legend.only) {
    if (!add) {
      par(plt = bigplot)
    }
    if (!info$poly.grid) {
      image(..., breaks = breaks, add = add, col = col)
    }
    else {
      poly.image(..., add = add, col = col, midpoint = midpoint, 
                 border = border, lwd.poly = lwd)
    }
    big.par <- par(no.readonly = TRUE)
  }
  if ((smallplot[2] < smallplot[1]) | (smallplot[4] < smallplot[3])) {
    par(old.par)
    stop("plot region too small to add legend\n")
  }
  ix <- 1:2
  iy <- breaks
  nBreaks <- length(breaks)
  midpoints <- (breaks[1:(nBreaks - 1)] + breaks[2:nBreaks])/2
  iz <- matrix(midpoints, nrow = 1, ncol = length(midpoints))
  if (verbose) {
    print(breaks)
    print(midpoints)
    print(ix)
    print(iy)
    print(iz)
    print(col)
  }
  par(new = TRUE, pty = "m", plt = smallplot, err = -1)
  if (!horizontal) {
    image(ix, iy, iz, xaxt = "n", yaxt = "n", xlab = "", 
          ylab = "", col = col, breaks = breaks)
  }
  else {
    image(iy, ix, t(iz), xaxt = "n", yaxt = "n", xlab = "", 
          ylab = "", col = col, breaks = breaks)
  }
  if (!is.null(lab.breaks)) {
    axis.args <- c(list(side = ifelse(horizontal, 1, 4), 
                        mgp = c(3, 1, 0), las = ifelse(horizontal, 0, 2), 
                        at = breaks, labels = lab.breaks), axis.args)
  }
  else {
    axis.args <- c(list(side = ifelse(horizontal, 1, 4), 
                        mgp = c(3, 1, 0), las = ifelse(horizontal, 0, 2)), 
                   axis.args)
  }
  
  if (!is.null(axis.at)) axis.args$at <- axis.at
  if (!is.null(axis.lab)) axis.args$labels <- axis.lab
  if (!is.null(axis.side)) axis.args$side <- axis.side
  
  do.call("axis", axis.args)
  box()
  if (!is.null(legend.lab)) {
    legend.args <- list(text = legend.lab, side = ifelse(horizontal, 
                                                         1, 4), line = legend.line, cex = legend.cex)
  }
  if (!is.null(legend.args)) {
    do.call(mtext, legend.args)
  }
  mfg.save <- par()$mfg
  if (graphics.reset | add) {
    par(old.par)
    par(mfg = mfg.save, new = FALSE)
    invisible()
  }
  else {
    par(big.par)
    par(plt = big.par$plt, xpd = FALSE)
    par(mfg = mfg.save, new = FALSE)
    invisible()
  }
}
```
