## Supplementary material 3 - R code for "A multidimensional framework for measuring biotic novelty: How novel is a community?": Simulation testing.R

```
# Testing BNI on simulated data

## Required packages: vegan, ape
source(file = "FUNCTION BNI.calc.R")
source(file = "FUNCTIONS BNI simulations.R")

## Scenario 1: random scenario with no trait difference between natives and neophytes
simRandom<- simul.BNI(simulation.trait = "random", nreps = 10)
plot.BNIsim(simRandom, type = c("Full"), color.option = "",color.bar = FALSE, diff.col.max = TRUE )

## Scenario 2:  effect of increasing trait differences between natives and aliens
simTraitDif <- simul.BNI(nreps = 10,maxDT = 10 )
plot.BNIsim(simTraitDif, type = "Full",  diff.col.max = FALSE )

## Scenario 3a: Test effect of increasing differences in trait variance 
simSDDif <- simul.BNI(simulation.trait = "variance difference",
                      maxDSD = 10,nreps = 10)
plot.BNIsim(simSDDif , type = "Full", color.option = "SD")

## Scenario 3b:  effect of increasing SD differences between natives and aliens if natives have already a high variance
simSDDif.natSD <- simul.BNI(simulation.trait = "variance difference", sim.SD = 5, maxDSD = 10, nreps = 10)
plot.BNIsim(simSDDif.natSD , type = "Full", color.option = "SD")

## Schenario 4a:  effect of increasing trait and SD differences between natives and aliens
simbothDif <- simul.BNI(simulation.trait = "both different", maxDSD = 5, nreps = 10)
plot.BNIsim(simbothDif , type = "Full",  color.option ="SD", which.bar = "together")

## Scenario 4b: effect of increasing trait and SD differences between natives and aliens, if negative covaration
simbothDif.inverse <- simul.BNI(simulation.trait = "both different", maxDSD = 5,maxDT = 10, sd.trend = "inverse", nreps = 10)
plot.BNIsim(simbothDif.inverse , type = "Full",   color.option ="mean", which.bar = "inverse")
 
```
