## Supplementary material 4 - Supplementary figures for "A multidimensional framework for measuring biotic novelty: How novel is a community?"

### Supplementary material S4 – Supplementary figures

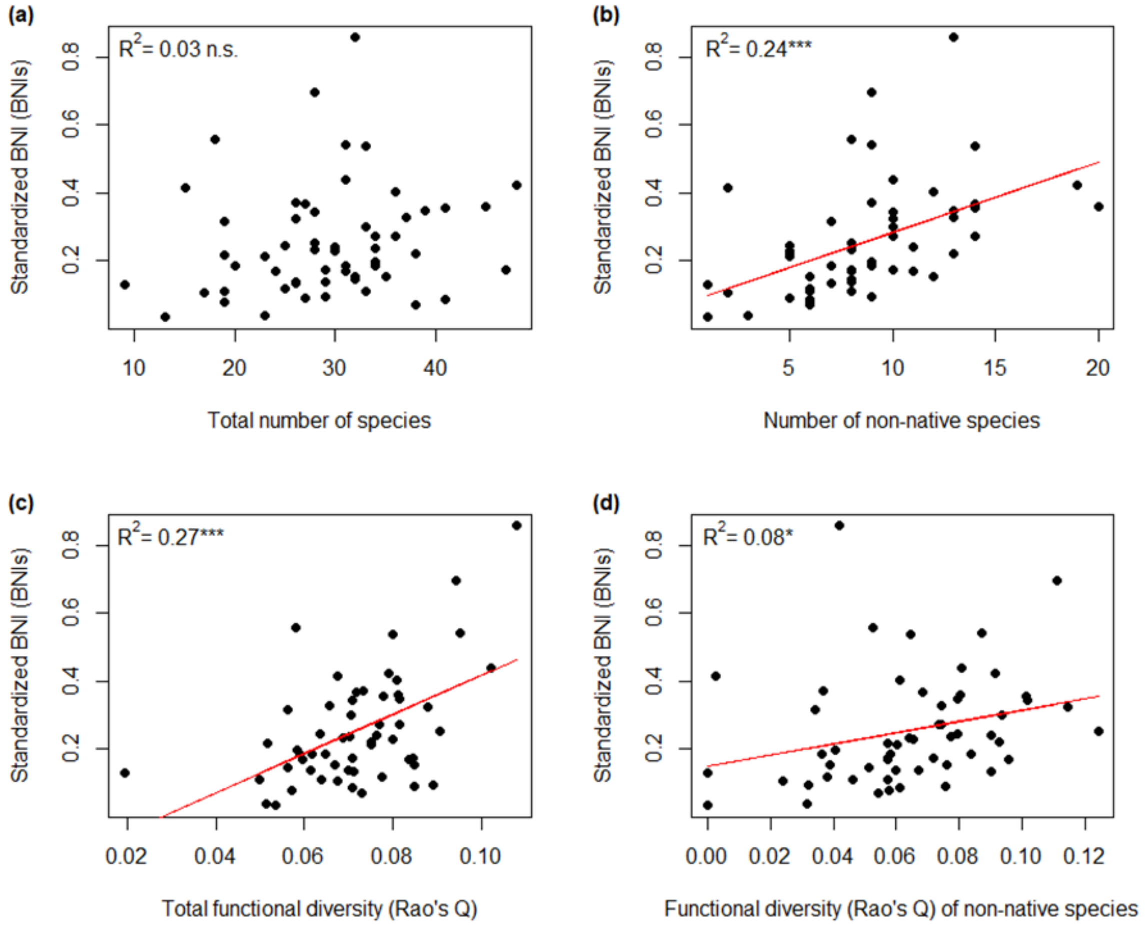


**Figure S4.1. Case study 1:** Relationships between the standardized BNI (BNIs) and (a) the total number of species, (b) the number of non-native species, (c) Rao's quadratic entropy as a measure of functional diversity, and (d) the functional diversity of non-native species in the 56 urban grassland plots. Asterisks indicate statistical significance using linear models ('***' = *P* < 0.001, '**' = *P* < 0.01, '*' = *P* < 0.05, 'n.s.' = *P* ≥ 0.05).

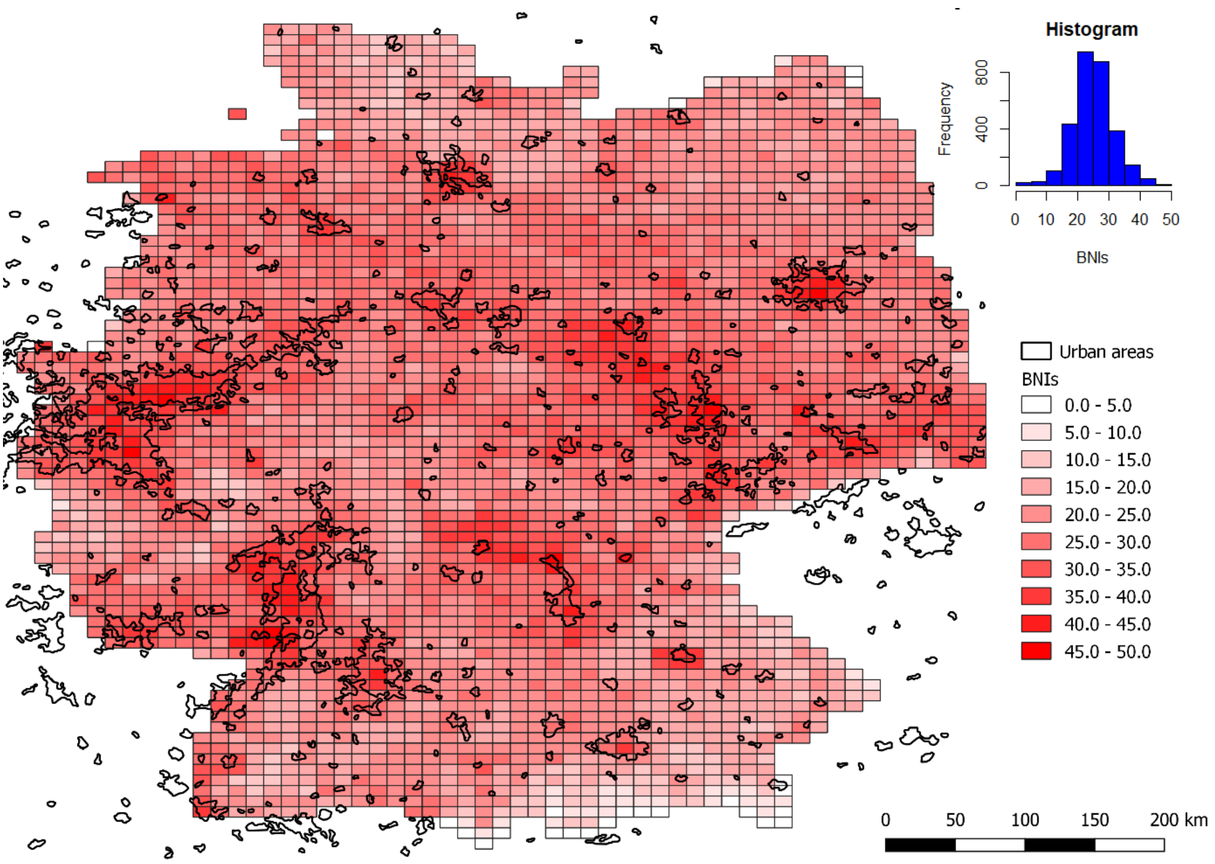


**Figure S4.2. Case study 2:** Biotic novelty of co-occurring vascular plants in Germany aggregated in 11 x 11 km grid cells calculated with the standardized version of the BNI (BNIs). Areas outlined in black indicate the extent of urban areas based on MODIS satellite data (Schneider *et al.* 2009).
